# High genetic diversity of mpox virus (MPXV) in three different rodent species in the Democratic Republic of the Congo (DRC)

**DOI:** 10.1101/2025.08.28.672325

**Authors:** Meris Matondo Kuamfumu, Pascal Baelo, Audrey Lacroix, Grace Odia Kadima, Simon-Pierre Ndimbo Kumugo, Eddy Kinganda Lusamaki, Antoine Nkuba Ndaye, Jill-Léa Ramassamy, Tine Cooreman, Mare Geraerts, Adrienne Amuri Aziza, Prince Akil, Fiston Cikaya, Nicolas Laurent, Rianne van Vreedendaal, Lea Fourchault, Lea Joffrin, Anne Laudisoit, Laurens Liesenborghs, Koen Vercauteren, Kevin Ariën, Tessa de Block, Daan Jansen, Casimir Nebesse, Steve Ngoy, Claude Mande, Ahidjo Ayouba, Eric Delaporte, Prince Kaleme, Luis Flores-Girón, Almudena Marí-Saéz, Herwig Leirs, Sophie Gryseels, Placide Mbala Kingebeni, Erik Verheyen, Guy-Crespin Gembu, Joachim Mariën, Martine Peeters, Steve Ahuka Mundeke

## Abstract

Altough zoonotic spillover events continue to drive human mpox outbreaks in the Democratic Republic of the Congo (DRC), the wildlife reservoir of mpox virus (MPXV) remains unkown. To address this gap, we screened samples from 2,701 wild mammals, mainly rodents (59.7%), bats (26.4%) and shrews (12.1%). Only six (0.2%) animals were Orthopoxvirus (OPV) PCR positive. Near full-length MPXV sequences were obtained from two squirrels (*Funisciurus anerythrus* and *Paraxerus* sp.) and one soft furred mouse (*Praomys jacksoni*). A novel Taterapox virus was identified in a shrew (*Crocidura cf. denti*). All newly identified MPXV strains belong to clade Ia, but they cluster into different groups or subgroups, despite being collected from geographically close locations, and all are closely related to human MPXV strainsfrom the same regions. Our study provides for the first time clear evidence that MPXV diversity is not restricted to a single rodent host species nor confined to a geographic area. Importantly, MPXV positive *Paraxerus* and *Praomys* specimens were sampled close to Kisangani, a city with more than one million inhabitants, highlighting that spillover events can als ooccur in or near major cities, with more favorable conditions for interhuman transmissions and potential emergence of new lineages.

## Introduction

Mpox, previously known as monkeypox, is a zoonotic disease caused by mpox virus (MPXV), of the *Orthopoxvirus* genus ^1^. The virus was first isolated in 1958 in Denmark from an imported macaque (*Macaca fascicularis*) ^2^. A decade later, the first human case was detected in a child with a smallpox-like disease in the Equateur Province of the Democratic Republic of the Congo (DRC) ^3^. Subsequent human cases were primarely reported from tropical rainforest areas of central and west Africa, where viral clades I and II circulate endemically, respectively ^1,4^. Initially, only sporadic zoonotic outbreaks were observed with limited person-to-person transmission ^5^. However, the eradication of smallpox and the cessation of smallpox vaccination led to increase of mpox incidence in endemic areas ^5–8^. This changed dramatically in 2022, when a clade II strain, namely subclade IIb related to an outbreak in Nigeria started to spread efficiently through sexual networks in countries classified as nonendemic, prompting WHO to declare mpox a Public Health Emergency of International Concern ^9,10^. Although the global incidence declined, this outbreak is still ongoing in 2025 with currently more than 100,000 cases recorded in 115 non endemic countries ^11^.

In the DRC, the number of mpox cases increased exponentially since 2020, with cases now reported in each province ^8^. Furthermore, during 2023-2024, two new lineages linked to sustained human-to-human transmission, including in sexual networks, emerged in the country: one in the east, designated clade Ib, and another in Kinshasa, representing a novel lineage within clade Ia ^12–15^. To identify the outbreaks with sustained human-to-human a new nomenclature system has been recently proposed, as such these outbreaks are now called clade Ib/sh2023 and clade Ia/sh2024 ^16^. The clade Ib/sh2023 outbreak has spread rapidely to other provinces in DRC and neighboring countries, leading the WHO to declare for the second time a Public Health Emergency of International Concern in August 2024 ^17^. Today, two transmission patterns contribute to human mpox cases in Africa: large human-to-human transmission chains in urban and densily populated areas and frequent zoonotic spillovers in rural endemic regions. ^18^. Since these spillovers drive the high genetic diversity of mpox, gaining a clearer understanding of the animal reservoir of the virus is essential for predicting, preventing, and controlling future outbreaks ^18^.

Altough African rodents are presumed to be the primary reservoir of MPXV, robust evidence on its natural hosts remains limited. Molecular identification or isolation of the virus is only available for two wild rodent species, including a Thomas’s rope squirrel (*Funisciurus anerythrus)* sampled in 1985 in the Mongala Province in northern DRC and a fire-footed rope squirrel (*Funisciurus pyrropus)* from Ivory Coast sampled in 2022 ^19,20^. Various other rodent species from different genera (*e.g., Cricetomys*, *Mastomys*, *Funisciurus, Graphiurus, Oenomys, Heliosciurus,*) and shrews (*e.g., Crocidura*) were also suggested as natural reservoirs ^21–26^. However, empirical evidence supporting these claims remains largely limited to serological data. Besides rodents, viral sequences have also been obtained from symptomatic sooty mangabeys (*Cercocebus atys*) and chimpanzees (*Pan troglodytes*) from Taï National Park in Ivory Coast as well as from a chimpanzee in a sanctuary in Cameroon^27–29^. However, due to their pronounced clinical symptoms, non-human primates are rather thought to be incidental hosts, like humans, that do not play a significant role in viral maintenance. To further document the role of rodents as reservoir species of MPXV, we screened wildlife samples collected across different regions of the DRC where MPXV is endemic.

## Results

### Animal species tested

Samples of a total of 2,701 animals that were captured at 31 different sites in nine provinces of the DRC (**Fig. 1**) were used in this study, representing 59.7% (n=1613) rodents, of which 587 squirrels; 26.4% (n=714) bats; 12.1% (n=327) shrews and 1.7% (n=47) animals belonging to other mammalian orders obtained opportunistically from local hunters or wild meat sellers. Most animals were captured in Tshopo Province (37.2%; n=1005), followed by Tshuapa (21.6%; n=584) and Bas-Uele Provinces (18.2%; n= 493). Squirrels were captured in 22 sites across seven provinces. The details of the different genera and species captured in each site and province are summarized in **Table 1** and **Supplementary Table S1**.

**Figure 1.**
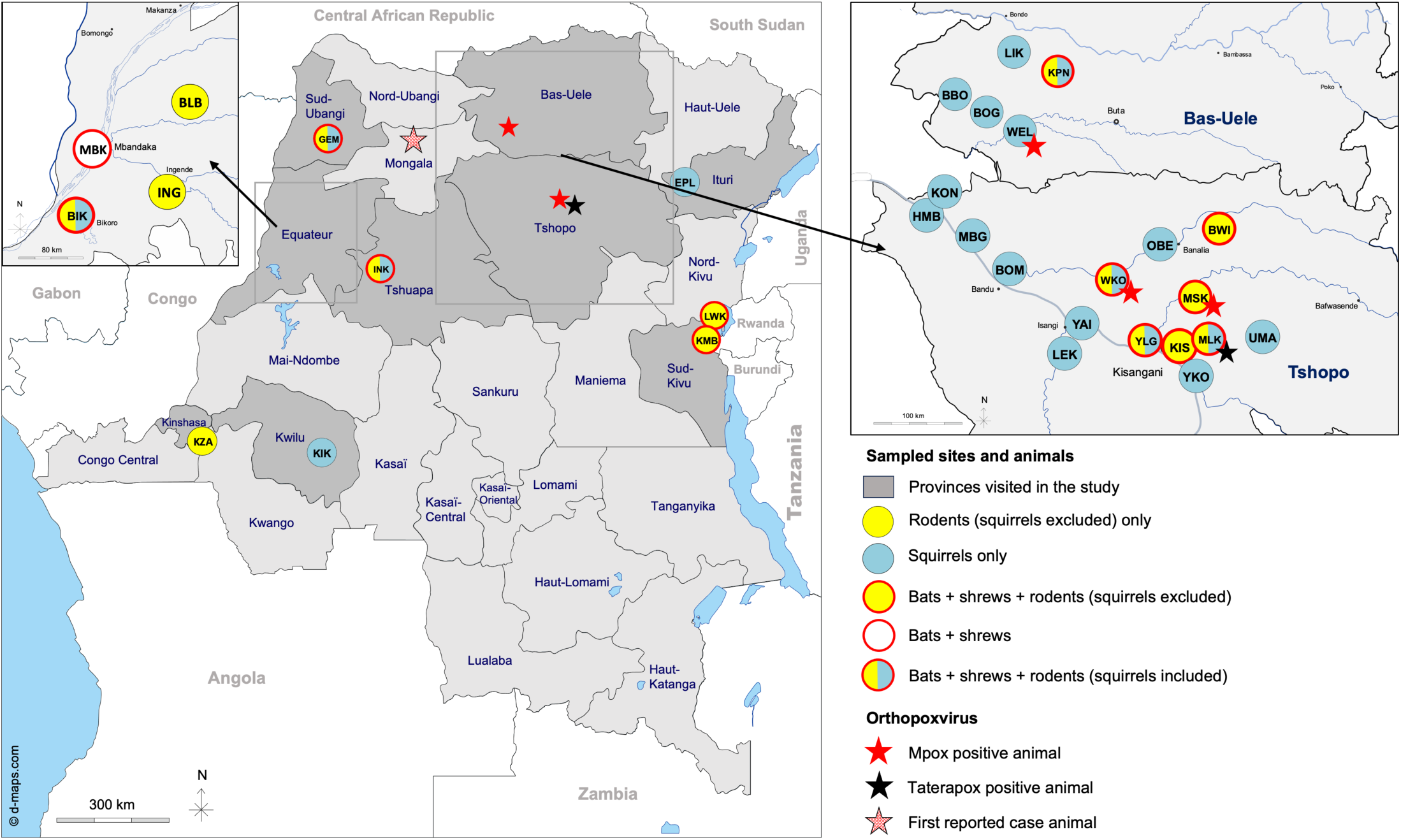
Small mammal capture sites. Animals have been captured between 2010 and 2024 in forests and scrublands in the Democratic Republic of Congo. Sites are highlighted with circles. Different colors are used to highlight the different animal orders captured on each site: yellow for rodents (excluding squirrels); yellow dot with red circle for bats, rodents and shrews; white dot with red circle for shrews and bats; dots with both yellow and blue and a red circle for bats, rodents, including squirrels and shrews; blue circle for squirrels only. Stars show sites where mpox (red) or taterapoxvirus (black) was detected in an animal. Sites are abbreviated with a three letter code as follows: Kimwenza (KZA), Mbandaka (MBK), Bikoro (BIK), Ingende (ING), Bolomba (BLB), Bulu (BUL), Gemena (GEM), Bawi (BWI), Kisangani (KIS), Yelenge (YLG), Maleke (MLK), Masako (MSK), Weko (WKO), the Lwiro-Kahungu axis (LWK), the Kavumu-Mbayo axis (KMB), Bogala (BOG), Bomane (BOM), Bombongolo (BBO), Epulu (EPL), Hembe (HMB), Inkanamongo (INK), Kikwit (KIK), Kona (KON), Kponyo (KPN), Likati (LIK), Mombongo (MBG), Obenge (OBE), Uma (UMA), Lieki (LEK), Wela (WEL), Yaikela (YAI) and Yoko (YKO).

**Table 1.**
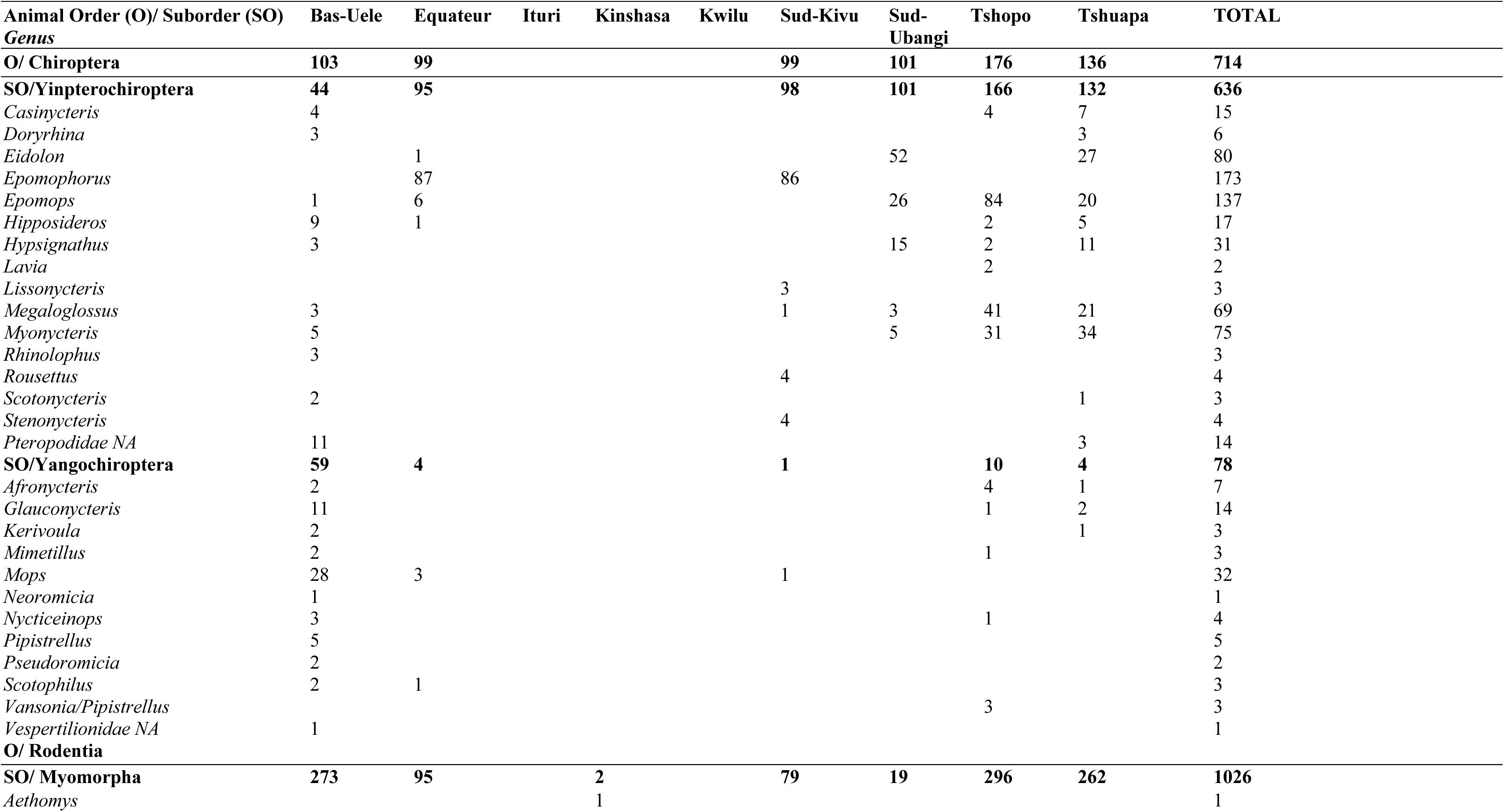
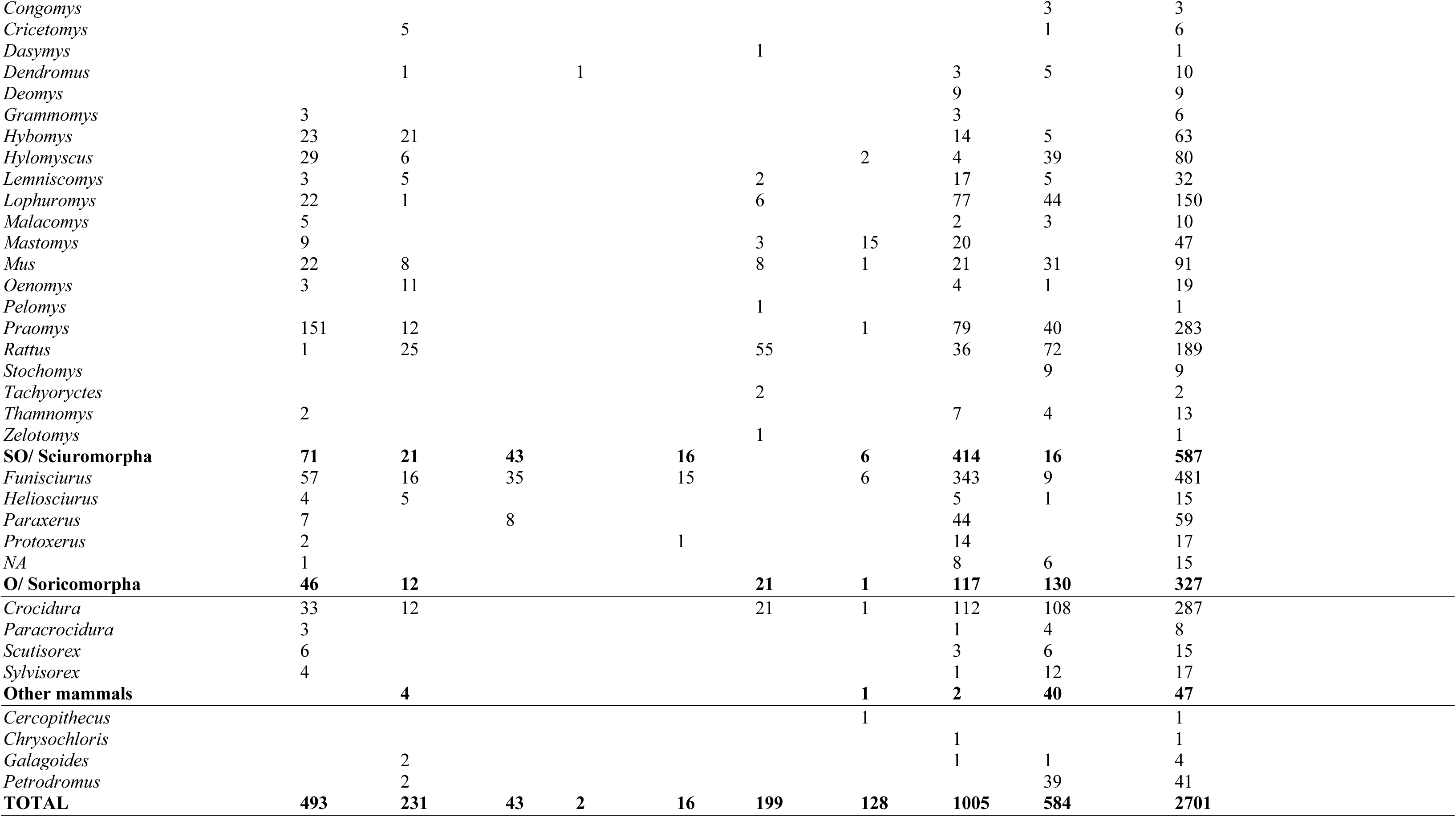
Animals sampled and tested by PCR for the presence of Orthopoxviruses in each province. Provinces correspond to those shown in **Fig. 1**. Data are shown at animal genus level. Animal species and collection sites per province are detailed in **Supplementary Table S1**.

Among the 714 bats, 636 and 78 were from the Yinpterochiroptera and Yangochiroptera suborder, respectively. The captured Yinpterochiroptera bats represent 16 different genera and at least 21 species. The most frequently captured genera were *Epomophorus* (27.2%, 173/636) and *Epomops* (21.5%, 137/636), followed by *Eidolon* (*i.e.*, *Eidolon helvum*) (16.6%, 80/636), *Myonycteris* (11.8%, 75/636) and *Megaloglossus* (10.8%, 69/636). At least 11 Yangochiroptera bat genera were captured, with *Mops* being the most frequent, 41.0% (32/78). Four different shrew genera from at least 16 species were tested, predominantly *Crocidura* (87.8%, 287/327). Squirrels from at least 5 genera were captured, mostly *Funisciurus* (81.9%; 481/587) and *Paraxerus* (10.1%; 59/587). For 15 squirrels the genus/species could not be identified. Among the other rodents, at least 22 genera from 45 species were captured, predominated by *Praomys* sp. (27.6%, 283/1,026), followed by *Rattus* sp. (18.4%, 189/1,026) and *Lophuromys* (*i.e.*, *Lophuromys dudui*) (14.6%, 150/1,026).

### PCR testing for presence of orthopox viruses

A total of 3,052 tissue samples from the 2,701 animals were analyzed for the presence of Orthopox viruses (OPV) (**Supplementary Table S2)**. Only six (0.2%) animals tested OPV PCR positive: four rodents among which three squirrels, and two shrews (**Table 2**). One of the rodents was a Jackson’s soft-furred mouse (PFPX-011/L818124) (*Praomys jacksoni*, Muridae family), captured in old fallow land in the Masako territory approximately 14 km from Kisangani (the capital of Tshopo Province). The three OPV positive squirrels belong to two different genera/species, *i.e.* one *Paraxerus* sp. (YLG044) captured in secondary forest near the town of Yelenge in Tshopo Province and two *Funisciurus anerythrus,* captured at Ahupa in Bas-Uele Province (UAC902) and at Yaekala in Tshopo Province (CRT57). Furthermore, two shrews tested positive; one specimen of *Crocidura cf. denti* (PFPX-175) collected from forested regions in the Maleke territory of Tshopo Province and one specimen of *Crocidura Olivieri* (RG002), captured along the Lwiro-Kahungu axis, Northeast of Bukavu (the provincial capital of Sud-Kivu Province), respectively (**Fig. 1**, **Table 2).**

**Table 2:**
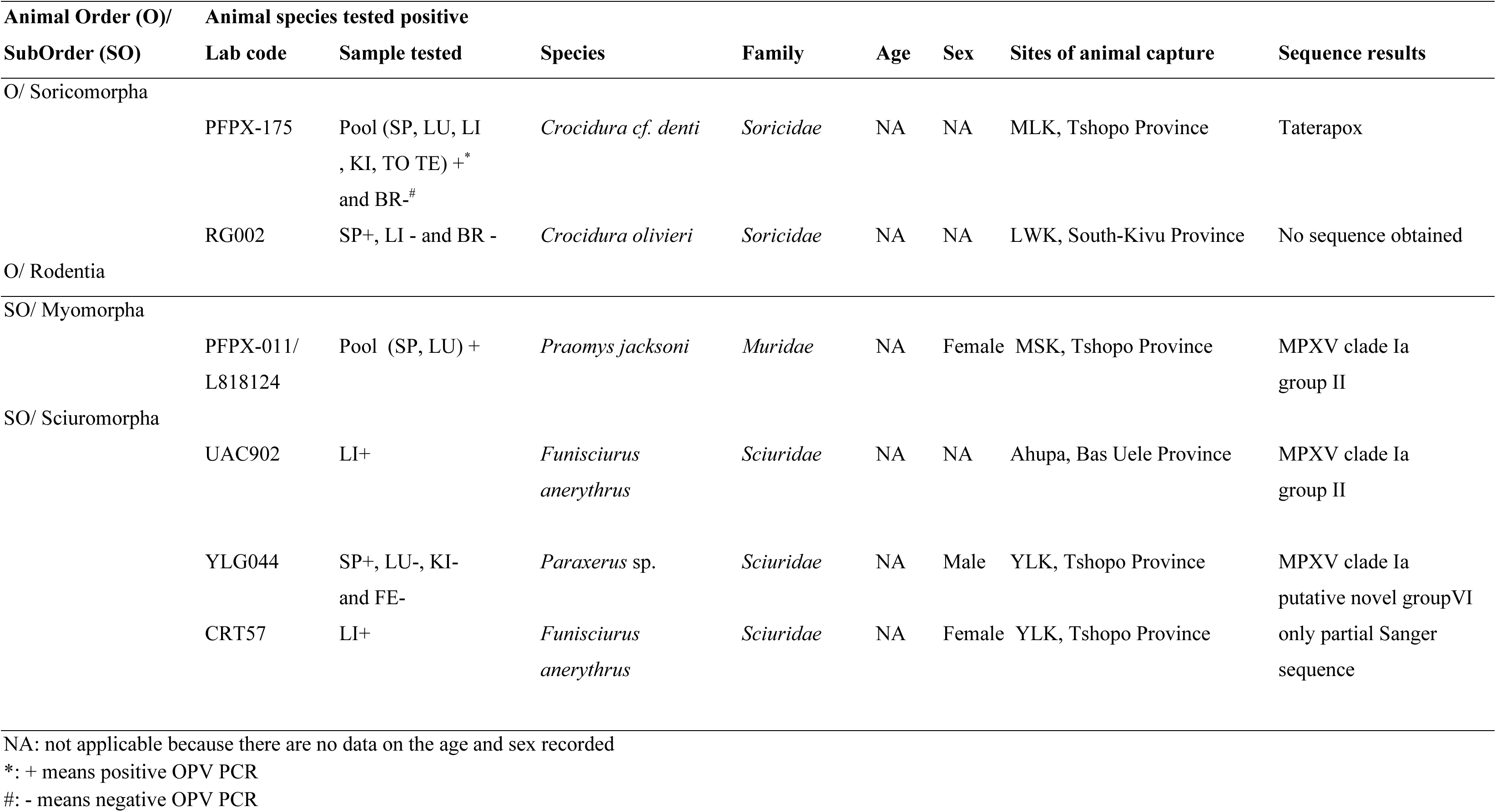
Details on the animals that tested positive using Orthopoxvirus PCR. Sample collection sites are represented by the same three-letter code as in **Fig. 1** (MLK: Maleke, LWK: Lwiro and Kahungu axis, MSK: Masako, YLK: Yelenge). Types of samples tested are listed using same abbreviations as in **Supplementary Table S2** (SP: Spleen, LI: Liver, LU: Lung, KI: Kidney, TO: Tongue, BR: Brain, TE: Testis and FE: Faeces).

### Sequencing and phylogenetic analysis of orthopox PCR positive samples

Whole genome sequencing confirmed the presence of MPXV clade Ia in *Praomys jacksoni*, *Paraxerus* sp. and one *Funisciurus anerythrus*. Near full-length sequences were obtained for the MPXV strains from the *Praomys jacksoni* (PFPX-011/L818124; 70% genome coverage, 9.6 mean sequence depth) and from the *Paraxerus* sp. (YLG044; 95% genome coverage, 1601 mean sequence depth). Full-length genome sequence was obtained for the MPXV from the *Funisciurus anerythrus* (UAC902; 99.8% genome coverage, 3072 mean sequence depth). Sanger sequencing of the OPXV positive PCR fragment confirmed OPXV for the *Funisciurus anerythrus* but not for the *Crocidura olivieri* captured in Tshopo and Sud-Kivu Provinces, respectively. Interestingly, sequence analysis from the *Crocidura cf. denti* revealed a near full length Taterapox virus genome (PFPX-175; 88.2% genome coverage and 96 mean sequence depth (**Table 2**, **Fig. 2**).

**Figure 2.**
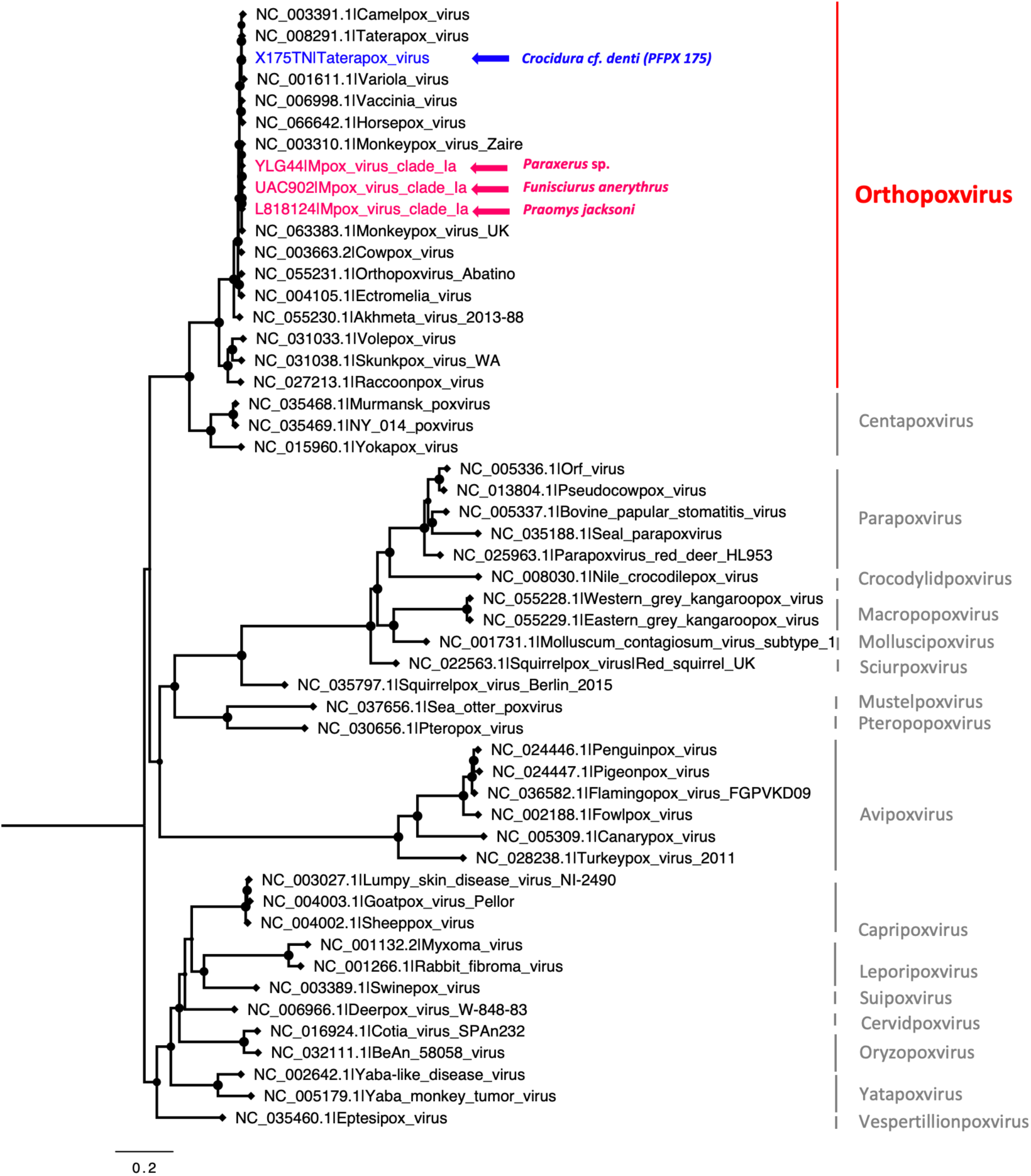
Phylogenetic tree with the newly generated near-complete MPXV and OPXV genomes. Newly obtained sequences were aligned with other chordopoxviruses reference genomes as described in Methods. The Maximum Likelihood tree was built using the TVM+R+F5 model. The new strains are highlighted in red for MPXV and blue for the new Taterapox virus (TPXV).

Detailed phylogenetic analysis of the MPXV sequences revealed clustering with various clade Ia groups or subgroups (**Fig. 3**). In contrast to the previously reported MPXV sequence from the *F. anerythrus* collected in 1985 in Mongolo Province, which belongs to group III, the new MPXV strain from the *F. anerythrus* squirrel collected in 2017 from Bas Uele belongs to group II and fell in a separate cluster with recent human MPXV strains from the same province and human MPXV strains from 2018 reported from a health zone nearby the site where the positive animal was captured (**Fig. 3 and Fig. 4 A-B**). The new MPXV sequence obtained from *Praomys jacksoni* also belonged to group II, although in a different and distanly related cluster together with human MPXV sequences from adjacent health zones to the site where the animal was captured (**Fig. 3 and Fig. 4 C-D**). The MPXV sequence from the *Paraxerus* sp. squirrel, captured at approximately 50 km from the MPXV positive *Praomys,* clusters within a potential novel group (putative group VI, to be confirmed by a nomenclature committee) that has been recently reported in clade Ia ^18^. Similarly, as for the above-mentioned animal MPXV strains, the MPXV from the *Paraxerus* squirrel is close to contemporary human MPXV strains from nearby health zones, but in this case the strain is also very close to a recent human MPXV from the same healthzone (**Fig. 3 and Fig. 4E-F**). Moreover, Fig. 4E, also clearly illustrates the cocirculation of different MPXV lineages in the same healthzone, reflecting the diversity of viral strains that circulate in animals.

**Figure 3.**
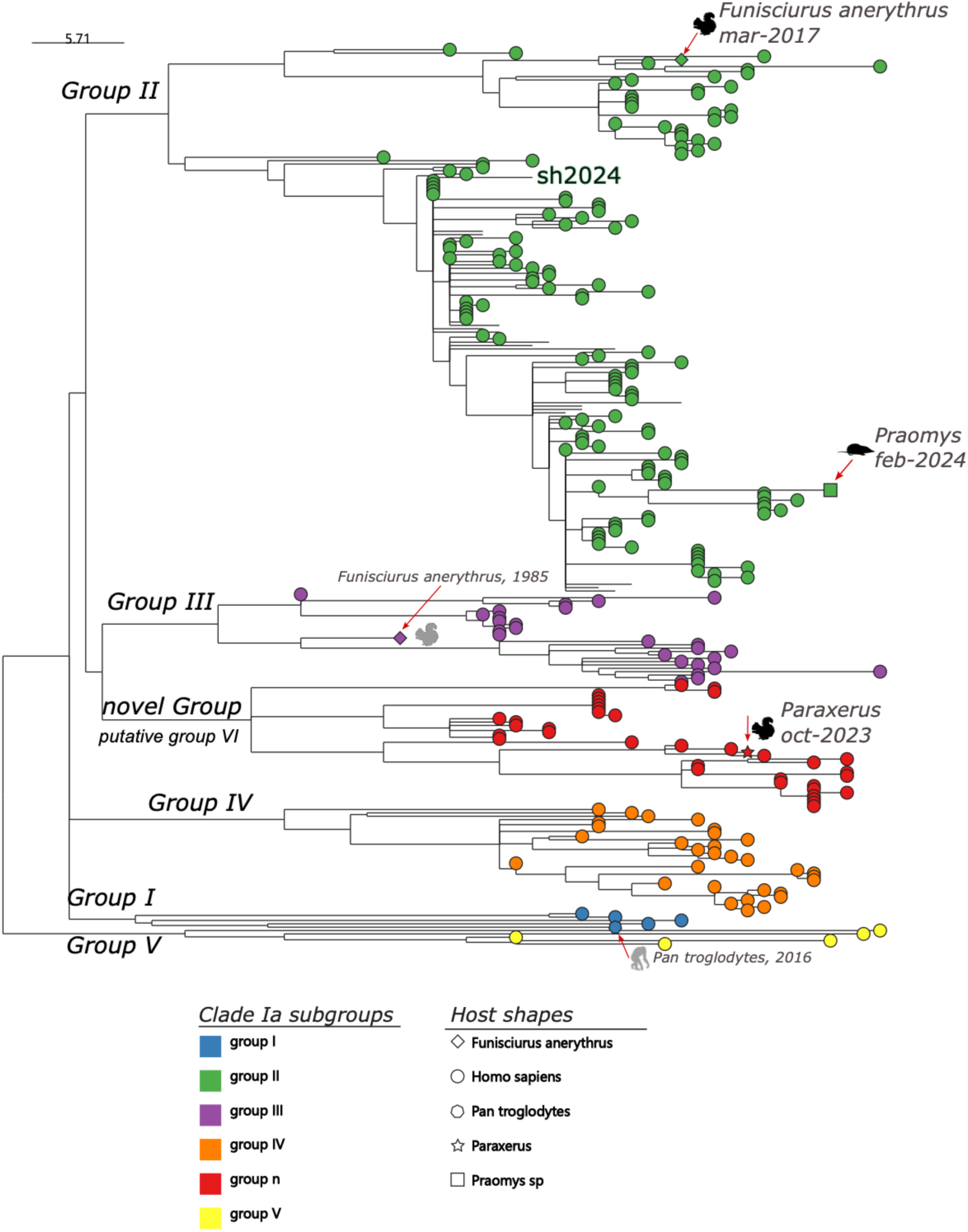
Phylogenetic tree with the newly obtained mpox virus (MPXV) strains from wild rodents captured in the Democratic Republic of Congo (DRC) aligned with other clade Ia reference genomes. The new sequences from animals are highlighted with a different symbol, their genus and/or species as well as collection date of the corresponding host species. The different groups within clade Ia are highlighted in different colors; novel group represents putative group VI. Previously published clade Ia sequences from a squirrel (*Funisciurus anerytrus*; DRC, 1985) and chimpanzee (*Pan troglodytes troglodytes*; Cameroon, 2016) are also included in the tree.

**Figure 4.**
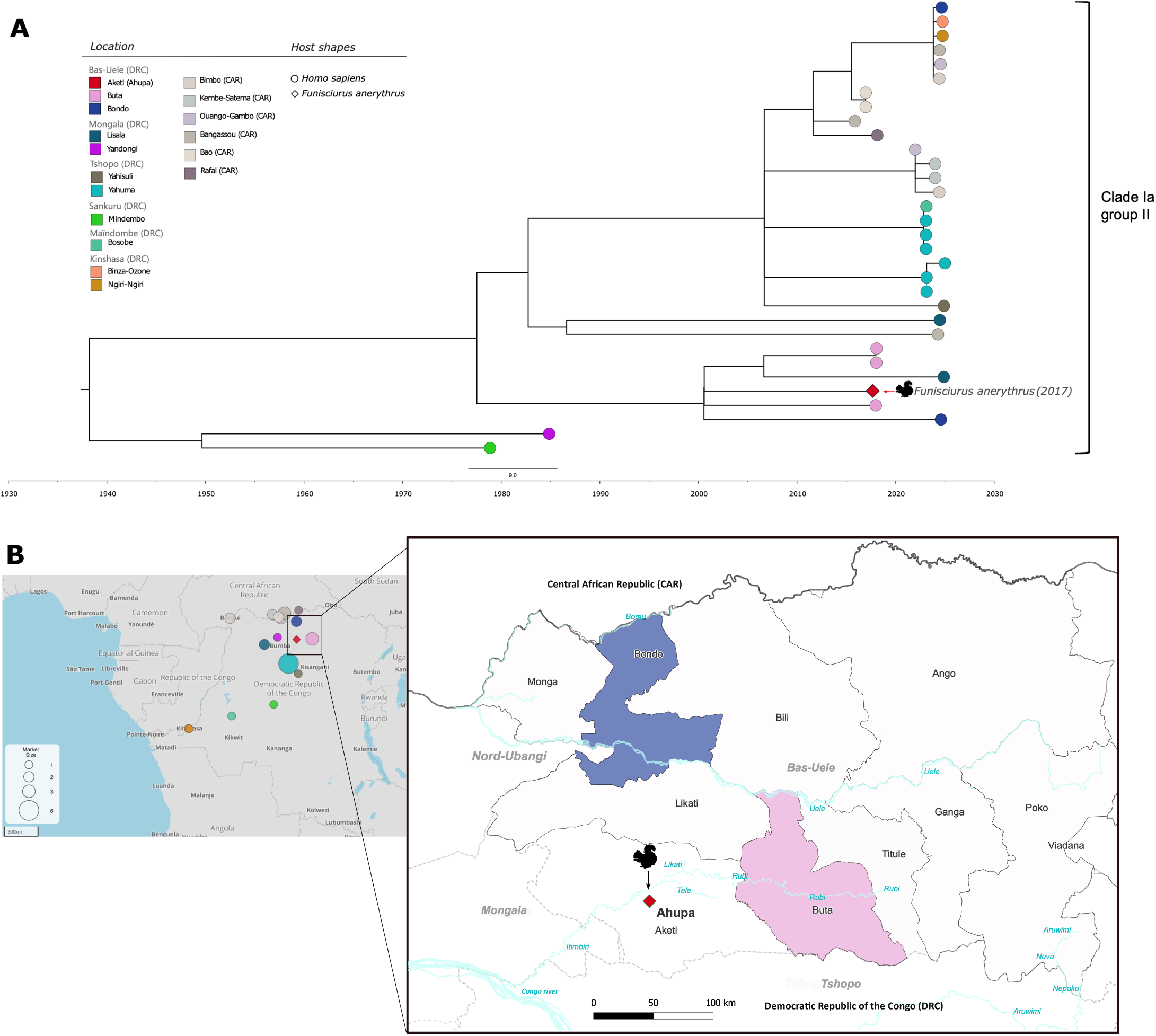
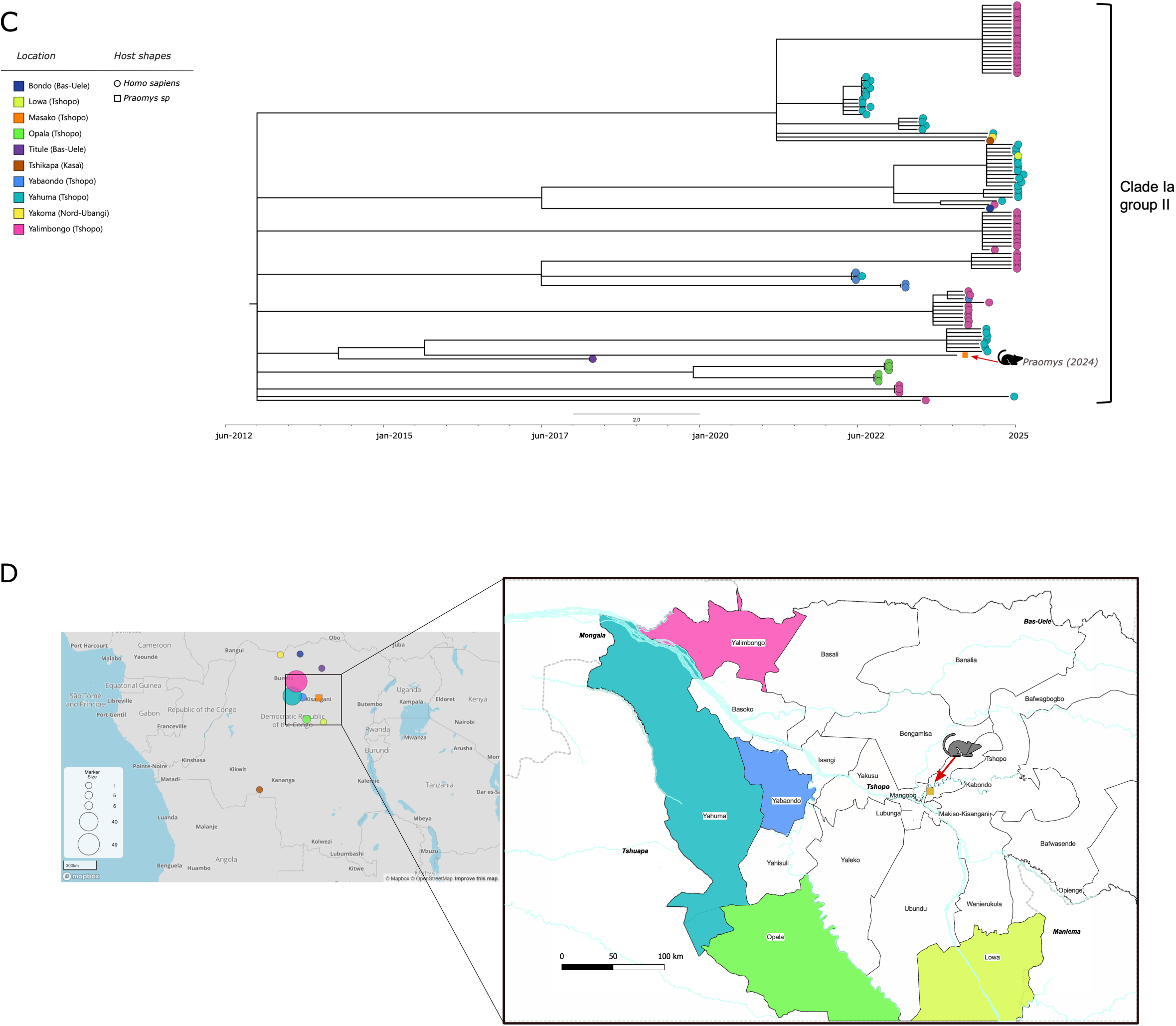
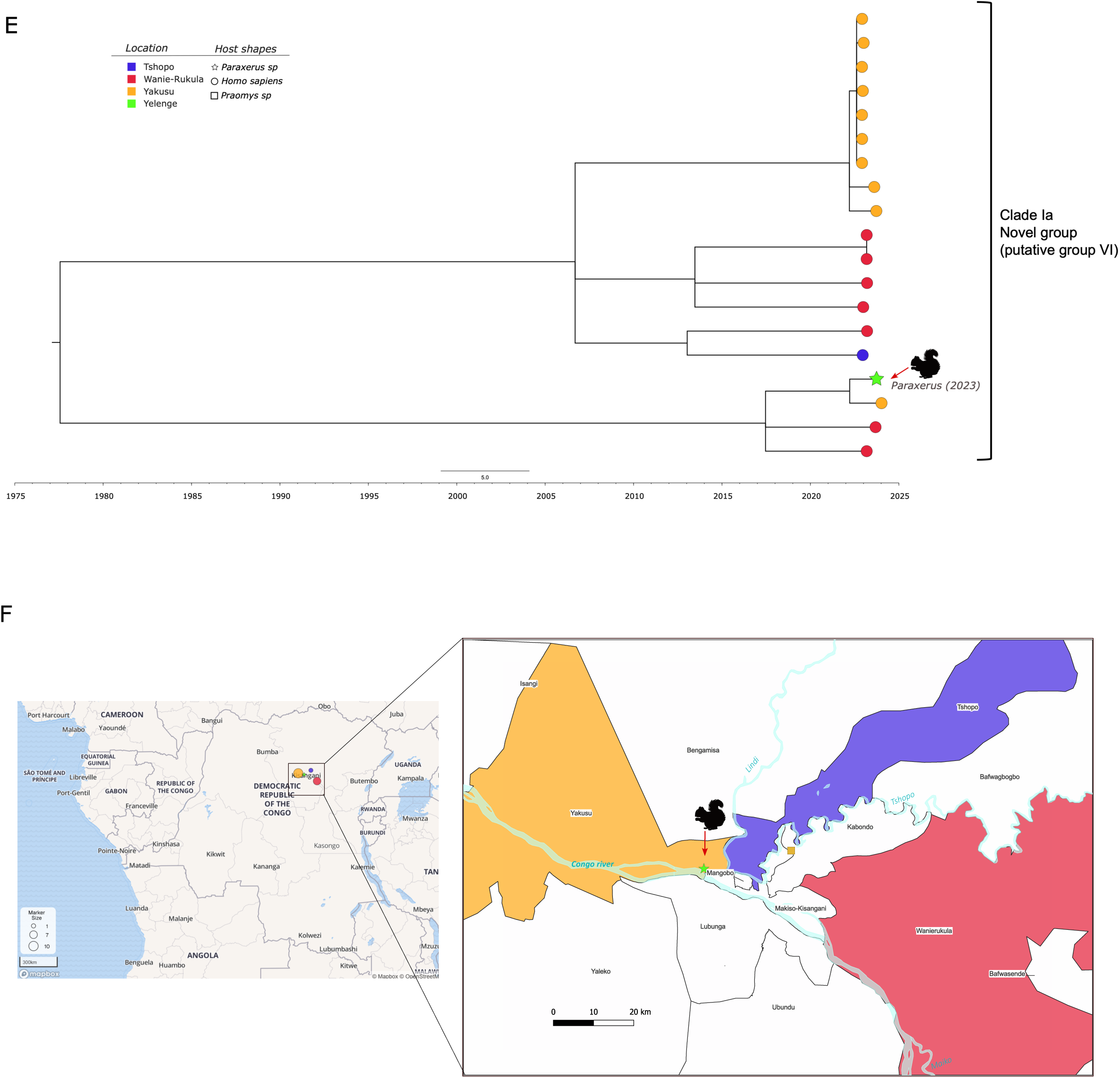
Phylogenetic sub-time trees highlighting new rodents MPXV strains and their respective most closely related human MPXV sequences; *Funisciurus anerythrus* collected in Bas-Uele province in 2017 (4A-B), *Praomys jacksonii* collected in 2023 in Tshopo Province (4C-D) and *Paraxerus* sp. collected in 2024 in Tshopo province (4E-F). For each MPXV sequence isolated from rodent, a phylogenetic sub-time tree highlights the most closely related human MPXV with tips colored according to the corresponding health zone (A, C, E). The geographic location of the animal and the different health zones centroids are shown on a map with circles in different colors and size of the circles reflecting the number of human MPXV sequences available for these health zones. Zoom panels display the geographic position of the host animal and the ranges of the health zones where most closely related human MPXV sequences were obtained (B, D, F).

### Detection of antibodies against MPXV

Samples from a subset of 1,356 bats and rodents (50.2% of the animals collected) for which dried blood spots (DBS) were available, were tested for antibodies against different MPXV virus antigens. For bats, 14/633 (2.2%) specimens tested positive against at least one antigen, including 6/80 (7.5%) of *Eidolon helvum*, 6/161 (3.7%) of *Epomophoru*s sp., 1/98 (1.0%) of *Epomop* sp. and 1/32 (3.1%) of *Mops* sp. (**Table 3 and Supplementary Table S3**). In rodents, 1.9% (14/723) individuals were positive for antibodies against at least one antigen, *i.e.* 7/215 (3.3%) of *Praomys* sp., 6/158 (3.8 %) of *Rattus* sp. and 1/72 (1.4%) from *Hylomyscus* sp. (**Table 3 and Supplementary Table S3**). Among all the animals tested, only two bats had antibodies against two MPXV antigens, and none had antibodies against all three.

**Table 3:**
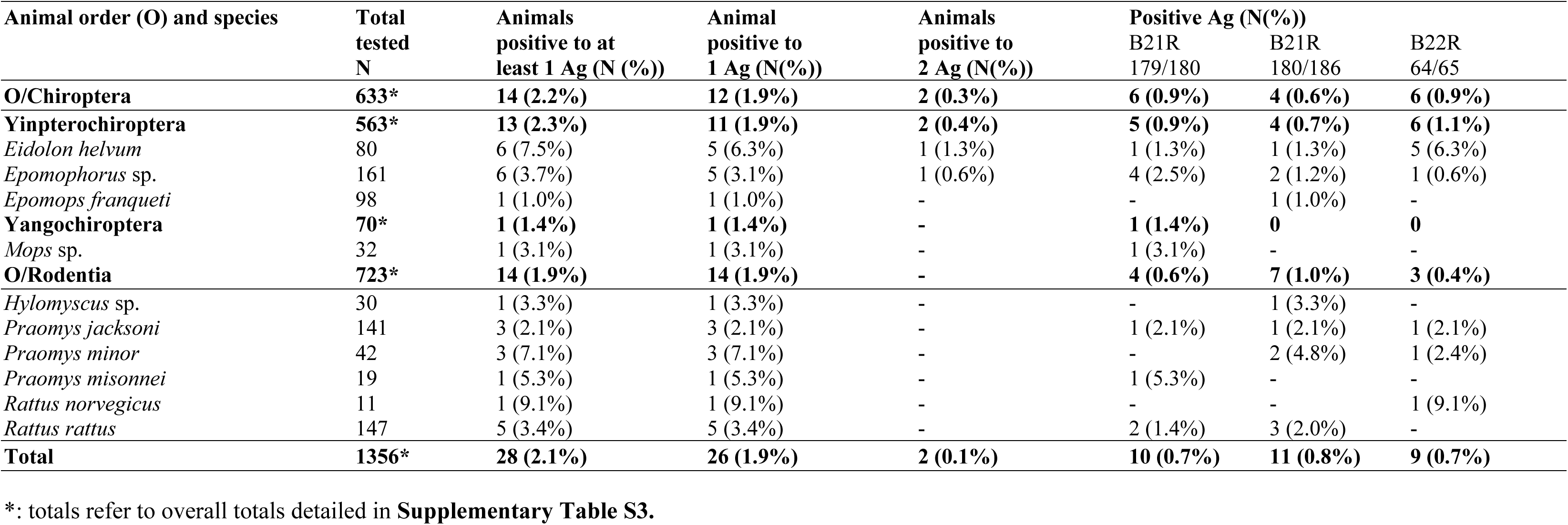
Number (and percentage) of animal species positive for presence of antibodies to at least one MPXV antigen by multiplex serology. Details on overall animals tested (including negatives) by species and animal order are displayed in **Supplementary Table S3**.

## Discussion

This study investigated the presence of MPXV virus in wild mammals in the Democratic Republic of the Congo (DRC), where zoonotic spillover events continue to fuel human MPXV outbreaks ^18^. On a total of 2,701 small mammals tested from nine provinces, we obtained MPXV genome sequences from three wild rodents, two squirrels (*Funisciurus anerythrus* and *Paraxerus* sp.) and a soft furred mouse (*Praomys jacksoni*). In addition, a novel Taterapox virus was identified in a shrew (*Crocidura cf. denti*).

Our findings indicate that MPXV circulates in several distantly related rodent species and shows high genetic diversity. Although all newly identified MPXV strains belong to clade Ia, they cluster into different groups or subgroups, despite being collected from geographically close locations. For example, the *Paraxerus* and *Praomys* specimens were collected only 50 km apart; yet the virus identified in *Paraxerus* belongs to a recently observed novel subgroup (putative group VI) within clade Ia ^18^, while the strain detected in *Praomys* falls within group II. Similarly, the novel MPXV detected from *Praomys* and *Funisciurus*, collected approximately 250 km apart, belong both to subgroup II, but to distinct, distantly related clusters. Surprisingly, the new MPXV detected in *F. anerythrus* from Bas-Uele Province did not cluster in group III-unlike the strain previously isolated from the same species in 1985 in Mongala Province. Therefor, our observations show that MPXV diversity is not strictly associated with a single rodent host species nor confined to geographic areas, as already suspected in our recent large-scale study on human MPXV strains ^18^.

Importantly, the newly identified rodent-derived MPXV strains are closely related to those previously or recently isolated from human cases in the same regions. This strongly suggests that these wildlife strains are at the origin of the human outbreaks ^18,30^. It is also striking that the positive *Paraxerus* and *Praomys* specimens from our study were respectively sampled at 15 and 50 km distance from Kisangani, the capital city of the Thsopo Province with more than one million habitants. This city is connected with other regions in DRC and other countries by roads or via the Congo River reaching Kinshasa, the capital city of DRC. This highlights that spillover of MPVX may not only occur in remote areas but also in or near major cities, with more favorable conditions for interhuman transmissions and potential emergence of new lineages associated with human-to-human transmission chains. Such a situation might have occurred during the outbreak with clade Ib/sh2023 in Sud-Kivu Province that started in a mining city and rapidly spread to other provinces in DRC and neighboring countries ^13,14^ or the clade Ia/sh2024 strain in Kinshasa ^15^.

Only few other surveys have been conducted to detect MPXV in rodents in the DRC. These studies sampled a wide diversity of species and were conducted in Mongala (Yambuku, n=383) and Tshuapa Provinces (Boende, n=281) and detected only one MPXV positive squirrel ^19,22,25^. Despite the low number of samples in which MPXV DNA was detected, our data and those from previous studies suggest that among rodents, squirrels are most likely a major MPXV reservoir. This observation is further supported by a recent report on an mpox outbreak among non-human primates (*i.e.* wild sooty mangabeys) in Taï National Park, Ivory Coast, which showed a link with consumption of infected fire-footed rope squirrel (*Funisciurus pyrropus*) ^20^. Additionally, molecular screening of 1,038 museum skin samples from *Funisciurus* sp. from Central Africa, revealed 93 (9.0%) animals that tested positive by real time PCR in skin samples. However, no sequence analysis was performed to confirm MPXV sequences, and these results have to be taken with caution given the high prevalence as compared to less than 2% in our study in wild animals in their natural environment ^31^. Moreover, antibodies against MPXV have previously been detected in *Funisciurus* species ^22^, and a recent ecological niche modelling study identified squirrels, and *F. anerythrus* in particular, as important potential reservoirs of MPXV ^21^. The low prevalence of MPXV DNA detection in our study is most likely related to the short viremic period. For exemple experimental infections in *F. anerythrus* have shown viral shedding for two weeks supporting this hypothesis ^32^. Moreover, viral peaks or viral shedding can be related to age of animals tested or pregnancy or lactaction status of females as observed for other viruses like filoviruses or coronaviruses ^33,34^.

Although we could not further investigate the role of squirrels as MPXV reservoirs through serology, since blood samples were not available, our serological results suggest MPXV circulation in rodent genera beyond squirrels, most notably *Praomys* and *Rattus* which is in line with the detection of MPXV and confirmed by sequencing in a soft-furred mouse (*Praomys jacksoni*). Serological evidence in rodents was also reported in previous studies in the DRC using a wide diversity of serological techniques like haemagglutinin-inhibition, ELISA or neutralising antibodies; antibodies to MPXV were detected in terrestrial rodents (*Cricetomys gambianus* and *Mastomys natalensis*) and shrews (*Crocidura hildegardeae*) ^21,23,24,26,29^. Overall, seroprevalence in our study is close to those observed in previous reports from Central Africa like Tshuapa Province in DRC (1.6%), Republic of Congo (1.9%) and Cameroon (1.2%), but higher seroprevalence rates (around 20%) have been reported in rodents from other provinces in DRC (Kwilu, Tshuapa, and Kongo Central, Sankuru) or other countries like Ghana (5.4% to 16%) or in Zambia (10.6%) ^23–26,29^. These discrepancies likely reflect small sample sizes, different species sampled and differences in serological methodologies used. We also detected MPXV antibodies in 2.2% of bats, particularly in specimens from *Eidolon, Epomophorus, Epomops* and *Mops* species. Although the serological data suggest that MPXV may have a broader host range, we cannot exclude that the detected antibodies are non-specific or cross-reactivite with other Orthopoxviruses or even unrelated pathogens.

Interestingly, we also confirmed the presence of a Tateropox virus, through near-full-length sequencing, in a shrew (*Crocidura cf. denti*) captured at Maleke, Tshopo Province. Taterapox virus (TATV) is generally considered to be the closest relative of variola virus (VARV) and was originally isolated from an apparently healthy and wild gerbil (*Tatera kempi* or *Gerbilliscus kempi*) captured in the North of Benin in 1968 ^35^. Like MPXV, this finding suggests that other orthopoxviruses may also have a broader geographic and host range than previously recognized. Studies on extent of Tateropox infections in wildlife and zoonotic spill-over events to humans are also warranted.

Taken together, our observations show that MPXV diversity is not strictly associated with a single rodent host species nor confined to geographic area. Our study provides the first clear evidence that besides *Funisciurus* squirrels, other rodents can also carry MPXV. The detection of MPXV in *Praomys jacksoni* is notable, as it is one of the most common small mammal species in the rainforests of central and eastern Africa ^36^. Serological data further suggest the circulation of MPXV in *Praomys* and even indicates a wider host range for MPXV beside rodents. Importantly, the newly identified MPXV strains in rodents are closely related to those isolated from human cases in the same regions. Further research is urgently needed to clarify MPXV transmission dynamics, identify reservoir species, and assess the nature and frequency of human–animal interactions. Such studies are essential to better understand and prevent the recurrent zoonotic spillover events occurring in the DRC and other endemic regions in Africa.

## Methods

### Study sites and animals

Between 2010 and December 2024, animals were captured in their natural habitat across 31 different sites in nine provinces of the DRC: Equateur, Tshopo, Sud-Ubangi, Kinshasa, Tshuapa, Kwilu, Bas-Uele, Ituri and Sud-Kivu (**Fig. 1**, **Table 1**). Most of these regions are endemic for mpox and have experienced outbreaks of other zoonotic diseases, including Ebola Virus Disease (EVD). The majority of animals captured were bats, shrews and rodents including squirrels. Terrestrial mammals, rodents and shrews were captured using Sherman traps, snap traps or pitfall traps placed in houses, adjacent fields, and the surrounding forest. Traditional traps were employed for arboreal squirrels. In addition, opportunitic samples were taken from mammals that were caught by local communities on local markets or when hunters arrived at the village. Rodent traps were set at sunrise and checked either in the evening or the following morning. Bats were captured using mist nets or harp traps at roosting and foraging sites in the forest, along rivers, and near houses at the same sites where terrestrial mammals were also sampled. Captured animals were humanely euthanised with isoflurane and necropsied *in situ* in a field laboratory under biosafety conditions. Tissue samples (mainly spleen, kidney, liver and lung) were stored in ethanol, RNA later (Ambion, Austin, TX, USA) or DNA/RNA shield (Zymo Research, Irvine, CA, USA). For a subset of animals, dried blood spots (DBS) as well as oral and rectal swabs were also collected. Samples were stored in the field at ambient temperature for maximum 3 weeks and subsequently at –20°C in the laboratory. Data on capture sites (GPS coordinates), capture method, morphology (body measurements, weight, color), sex, age class (adult, juvenile) and visual identification of species were recorded in the field.

### DNA extraction

Molecular analyses were conducted either at the National Institute for Biomedical Research (INRB) in DRC or at the Institute of Tropical Medicine (ITM), the University of Antwerp Antwerp or the Royal Belgian Institute of Natural Sciences in Belgium. Depending on the laboratory, DNA was extracted from spleen, liver or kidney samples (or if not available, lung, or heart or swabs) using one of the following methods: NucliSens MiniMAG® system (Biomerieux, Marcy-l’Etoile, France), the Nucleospin Tissue mini kit (Machery-Nagel, Düren, Germany) or the QIAamp 96 DNA QIAamp HT kit (Machery-Nagel, Düren, Germany).

### Molecular confirmation of the animal species captured

To confirm the species identifications in the field, cytochrome b (Cyt b) PCR was performed on DNA extractions from organs tested for presence of mpox virus. The specific primers used were Cyt b-L140217 (forward) and Cyt b-H15905 (reverse) for bats, and Cyt b-L14723 (forward) and Cyt b-H15905 or Cyt b-H15915 (reverse) for rodents ^37–39^. In addition, certain species identifications were verified using 12S PCR targeting the 450 bp fragment of mitochondrial DNA with primers 12S-L1091 and 12S-H1478 ^40^. After Sanger sequencing, the new sequences were compared with known sequences available in GenBank (https://blast.ncbi.nlm.nih.gov/Blast.cgi) and the African Mammalia database (http://projects.biodiversity.be/africanmammalia). For sequences with no or low similarity (<97%) hits with species sequences available in Genbank, a phylogenetic tree was constructed using maximum likelihood methods implemented in PhyML with reference sequences from the closest genus/species identified by Blast analysis, to obtain at least genus identification using Seaview Version 5.0.4 ^39,41^. For certain animals, identification was only possible at the genus level, mostly due to the lack of reference sequences in Genbank.

### Molecular detection of viral DNA

At INRB, PCR screening was performed with an Orthopoxvirus PCR (OPV PCR), targeting a 269 bp fragment of the haemagglutinin (HA) gene, using primers OPV.HA-156F and OPV.HA-424R as previously described (Patrono et al, 2020). Briefly, DNA was amplified using the GoTaq Hot Start Master Mix PCR kit (Promega, Madison, WI, USA) as follows: Taq activation at 94°C for 2 min, 10 cycles of 92 ◦C for 30 s, 55 ◦C for 30 s with –0.5 ◦C/cycle and 72 ◦C for 30 s, 35 cycles of 92 ◦C for 30 s, 55 ◦C for 30 s and 72 ◦C for 30 s, with a final elongation at 72°C for 5 min. At ITM, a TaqMan Real-time PCR targeting the Orthopox virus (OPXV) P4A gene of all orthopoxviruses was performed ^28,42^ and positives cycle threshold (CT < 35) were confirmed using the above described conventional OPV. After agarose gel electrophoresis, PCR products of the expected size were sequenced by Sanger technology.

### Orthopox virus DNA sequencing

Whole genome sequencing was attempted on samples that were positive for OPXV PCR. Diversified approaches were used on different NGS platforms including either hybridization capture protocols (Custom Mpox Research Panel or Comprehensive Viral Research Panel; Twist Biosciences, California, USA), or amplicon-based protocols using either Yale or artic-inrb-v.1.0.1 primers ^15,18^. Final libraries were loaded onto Illumina (Iseq100 or Miseq, Illumina Inc., San Diego, CA, USA) or Nanopore (MinION or GridION; Oxford Nanopore Technologie (ONT), Oxford, UK) sequencers. Illumina raw data (FASTQ files) were processed and cleaned using GeVarLi pipeline (https://forge.ird.fr/transvihmi/nfernandez/GeVarLi) and analyzed subsequently by CZID (https://czid.org/), according to the primers scheme. ONT raw data (FAST5 or POD5 files) were basecalled and obtained FASTQ files were then processed for consensus genome generation using either the Metatropics pipeline for hybridization capture protocol or ARTIC-MPXV-nf for amplicon sequencing approaches (https://github.com/artic-network/artic-mpxv-nf). Clade assignment was performed using Nextclade (https://clades.nextstrain.org/).

### Orthopoxvirus phylogenetic analysis

Using MAFFT-7.505 ^43^, the newly generated near-complete MPXV and OPXV genomes were aligned with other chordopoxviruses reference genomes and one entopomopoxvirus as outgroup accessed in the NCBI GenBank on 22^nd^ April 2025. Best fit model based on Bayesian Information Criterion (BIC) was found using using Model finder implemented in IQ-TREE-2 multicore version 2.2.6 when running the ML phylogenetic analysis ^44–46^. Trees formatting and annotation was performed using FigTree (http://tree.bio.ed.ac.uk/software/Figtree/) and Inkscape.

### Mpox virus (MPXV) phylogenetic analysis

Phylogenetic analysis for MPXV was performed using nextstrain bioinformatic and visualisation workflow which included (i) Nextclade for pairwise sequence alignment using NC003310 reference, (ii) masking several regions of the genome, including the first 800 and last 6422 base pairs and multiple repetitive regions of variable length, (iii) Maximum Likelihood (ML) phylogenetic reconstruction using GTR model with IQTREE-2 ^45^ and (iv) ancestral state reconstruction and temporal inference using TreeTime ^47^. Trees were formatted and annotated using FigTree (http://tree.bio.ed.ac.uk/software/Figtree/), Microreact (https://microreact.org) and Inkscape.

### Screening for MPXV antibodies

Dried blood spots (DBS) were available for a subset of bat and rodent samples, and analyzed using our previously described multiplex MPXV serology with Luminex® technology ^48^. Briefly, three previously identified synthetic 31-mer peptides from right terminal region B (B21R.179/180 and B21R.185/186 for MPXV and B22R.64/65 for variola major) were synthesized with an additional N-terminal cysteine residue (82–95% purity) and conjugated to bovine serum albumin (BSA) (Eurogentec, Seraing, Belgium) ^48,49^. Conjugated peptides were covalently coupled on carboxyl functionalized fluorescent magnetic beads (5 μg/1.25 × 106) (Luminex Corp., Austin, TX, USA) with the BioPlex amine coupling kit (Bio-Rad Laboratories, Marnes-la-Coquette, France) according to the manufacturer’s instructions. After elution of the DBS as previously described ^50^, 100 µl of sample (1/200 final dilution) was incubated with 50 µl of beads (2,000 beads/μL in assay buffer) in 96-well flat-bottom plates (Greiner Bio-One, Frickenhausen, Germany). After washing, 50 µl of biotinylated anti-bat IgG (Bethyl, Montgomery, TX, USA) or anti rodent IgG (Sigma-Aldrich, St. Louis, USA) was added to each well and incubated in obscurity for 30 min. After washing, 50 µl of streptavidin-R-phycoerythrin (1 mg/ml) (Fisher Scientific/Life Technologies, Illkirch, France) was added in each well and incubated for 10 minutes under agitation (400 rpm/min) at room temperature. Antigen-antibody reactions were read on the MagPix (Luminex, Austin, TX, USA) or BioPlex-200 (BioRad, Marnes-la-Coquette, France). At least 50 events were read for each bead and expressed as Median Fluorescence Intensity (MFI) values. In the absence of positive controls for bats and rodents, we applied three different statistical methods to calculate the cut-off from the MFI values of the entire number of blood samples tested, as done in previous studies on seroprevalence of arboviruses or Ebola viruses in wildlife, *i.e.* change point analysis method and fitted univariate binomial and exponential distributions to define the cut-off at a 0.001 risk for error ^39,51^. Analyses were performed with R version 4.0.2 software (https://www.r-project.org). We then compared the cutoff values identified by the 3 different methods and calculated their mean as a consensus cutoff that we used in this study. Similarly, as in previous studies, we defined MPXV antibody positivity as reactivity to at least one peptide (*i.e.* MFI above cut-off value).

## Funding

This study received cofunding from PANAFPOX (funded by ANRS-MIE), PREACTS-ASAMCO (funded by the Agence Française de Développement (AFD)), RESOHLABO (Financed by Agence Francaise de Développement (AFD)), ZOOSURSY (funded by the EU Global Gateway, Project CA N° 700002203) projects; Research Foundation – Flanders (FWO, grant number G096222N) and 2018-2019 BiodivERsA joint call for research proposals under 481 the BiodivERsA3 ERA-Net COFUND program (grant number ANR-19-EBI3-0004). ITM’s SOFI (Structurele OnderzoeksFInanciering) programme supported by the Flemish Government, Science & Innovation and the DGD FA5 Synergy Fund. The research also received funding by the VLIR-UOS South Initiative project ‘Strengthening Academic Capacity to Respond and React to Monkeypox Epidemics: Discrimination and Origin of Eruptive Fevers in the Democratic Republic of Congo (DRC)’; BELSPO SSD project’ Congo Basin Integrated Monitoring For Forest Carbon Mitigation And Biodiversity’ (COBIMFO).

## Acknowledgements

CZID for providing the primers for sequencing analysis. The authors acknowledge the ISO 9001 certified IRD i-Trop HPC (member of the South Green Platform) at IRD montpellier for providing HPC resources that have contributed to the research results reported within this paper (https://bioinfo.ird.fr http://www.southgreen.fr).

## Institutional Review Board Statement

The study was reviewed by the Ethics Committee of the School of Public Health at the University of Kinshasa and approved for scientific publication under reference number ESP/CE/235/2024. For a part of the samples, there is no IACUC linked to the sampling because there was no dedicated animal ethical committee in the DRC at the time of sampling. The research permits and authorizations were granted by the University of Kisangani Biodiversity Surveillance Center (CSB [Centre de Surveillance de la Biodiversité] – in French) who has the scientific authority to do so. All “movement of personal” (called “ordre de Mission”) were signed and approved by the local authorities at each field site. Material transfer agreements (MTA) were issued by the University of Kisangani, Biodiversity Surveillance Center (CSB [Centre de Surveillance de la Biodiversité] – in French). All animals were captured using live traps and humanely euthanized (isofluorane) following the 2013 AVMA Guidelines for the Euthanasia of Animals and Sikes and Gannon 2007 (J Mammal. 88:809–23). Nagoya permits were obtained from all samples collected after 2014 that were sent to Belgium.

## Data and code availability

New mpox sequences generated during this study from mammals are deposited at Pathoplexus and GenBank repositories (Accession Numbers pending). The Taterapox virus genome has been submitted to GenBank (Accession Number pending). All the alignments and outputs files generated for the phylogenetic analysis and any additional information can be made available for research purposes from the lead contacts upon request. Scripts required to reproduce the analyses of this study can be accessed through GitHub at https://forge.ird.fr/transvihmi/nfernandez/GeVarLi for GeVarLi pipeline, https://github.com/artic-network/artic-mpxv-nf for ARTIC-MPXV-nf for ONT amplicon sequencing analyses, and through https://czid.org/ for CZID analyses.

## Conflicts of Interest

The authors declare no conflicts of interest.

## Author contributions

MP, SAM, PB, AL, AA, HL, EV, GCG, JM conceptualized the study and designed the methodology; SPNK, MMK, JLR, LJ, PB, PK, CM, SN, CN, LFG, GCG and AnL conducted field work; GKO, MMK, AL, MG, TC, NL, RvV, LJ, LF, TdB, PA, FC, AAA conducted laboratory work ; MMK, AL, EKL, PA, KV, TC, JM, EV and MP curated data of the field and/or laboratory work ; MP, AA, AMS, ANN, SAM, PMK, HL, SG, EV, GG, JM, LFG supervised laboratory and/or field activities ; MMK, PB, MP, JM, AL, EKL wrote the initial draft ; MP, AA, ED, SAM, LL, KA, HL, EV, JM, GCG obtained funding for the study; All authors reviewed the document and approved the final manuscript.

## Supplementary Information

**Supplementary Table S1.** Animals sampled and tested by PCR for the presence of Orthopoxviruses per province and collection site, by animal type, genus and species. Provinces correspond to those shown in Fig. 1.

**Supplementary Table S2.** Number of samples (N) analyzed by PCR for the presence of Orthopoxvirus by sample type. Sample types are listed using two-letter abbreviation, and organ pools are designated from pool 1 to 4. Sample number testing positive for OPV PCR are into brackets. Tested organs tested are identified as follow: SP: Spleen, LI: Liver, LU: Lung, KI: Kidney, FE: Faeces, IN: Intestine, SA: Oral swab, RE: Rectal swab, TE: Testis, TO: Tongue, EY: Eye, BR: Brain. Pools contain the following organs: Pool 1: SP+LU; Pool 2: LI+KI; Pool 3: SP+LI+LU+KI+TE+TO and Pool 4: TE+BR.

**Supplementary Table S3:** Number of animals tested for presence of antibodies against MPXV antigen by Luminex serology, by animal species and order.

## Notes

### Competing Interest Statement

The authors have declared no competing interest.

## References

1. Ulaeto, D. et al. New nomenclature for mpox (monkeypox) and monkeypox virus clades. Lancet Infect Dis 23, 273–275 (2023).

2. von Magnus, P., Andersen, E. K., Petersen, K. B. & Birch-Andersen, A. A Pox-Like Disease in Cynomolgus Monkeys. Acta Pathologica Microbiologica Scandinavica 46, 156– 176 (1959).

3. Ladnyj, I. D., Ziegler, P. & Kima, E. A human infection caused by monkeypox virus in Basankusu Territory, Democratic Republic of the Congo. Bull World Health Organ 46, 593– 597 (1972).

4. Arita, I., Jezek, Z., Khodakevich, L. & Ruti, K. Human monkeypox: a newly emerged orthopoxvirus zoonosis in the tropical rain forests of Africa. Am J Trop Med Hyg 34, 781–789 (1985).

5. Gessain, A., Nakoune, E. & Yazdanpanah, Y. Monkeypox. New England Journal of Medicine 387, 1783–1793 (2022).

6. Rimoin, A. W. et al. Major increase in human monkeypox incidence 30 years after smallpox vaccination campaigns cease in the Democratic Republic of Congo. Proc Natl Acad Sci U S A 107, 16262–16267 (2010).

7. Sklenovská, N. & Van Ranst, M. Emergence of Monkeypox as the Most Important Orthopoxvirus Infection in Humans. Front Public Health 6, 241 (2018).

8. Bangwen, E. et al. Suspected and confirmed mpox cases in DR Congo: a retrospective analysis of national epidemiological and laboratory surveillance data, 2010–23. The Lancet 405, 408–419 (2025).

9. WHO. WHO Director-General declares the ongoing monkeypox outbreak a Public Health Emergency of International Concern. https://www.who.int/europe/news/item/23-07-2022-who-director-general-declares-the-ongoing-monkeypox-outbreak-a-public-health-event-of-international-concern (2022).

10. O’Toole, Á. et al. APOBEC3 deaminase editing in mpox virus as evidence for sustained human transmission since at least 2016. Science 382, 595–600 (2023).

11. CDC. Mpox in the United States and Around the World: Current Situation. Mpox https://www.cdc.gov/mpox/situation-summary/index.html (2025).

12. Kibungu, E. M. et al. Clade I–Associated Mpox Cases Associated with Sexual Contact, the Democratic Republic of the Congo. Emerg Infect Dis 30, 172–176 (2024).

13. Masirika, L. M. et al. Ongoing mpox outbreak in Kamituga, South Kivu province, associated with monkeypox virus of a novel Clade I sub-lineage, Democratic Republic of the Congo, 2024. Euro Surveill 29, 2400106 (2024).

14. Vakaniaki, E. H. et al. Sustained human outbreak of a new MPXV clade I lineage in eastern Democratic Republic of the Congo. Nat Med 30, 2791–2795 (2024).

15. Wawina-Bokalanga, T. et al. Epidemiology and phylogenomic characterisation of two distinct mpox outbreaks in Kinshasa, DR Congo, involving a new subclade Ia lineage: a retrospective, observational study. The Lancet 406, 63–75 (2025).

16. Ruis, C. et al. A systematic nomenclature for mpox viruses causing outbreaks with sustained human-to-human transmission. Nature Medicine (2025) doi:10.1038/s41591-025-03820-6.

17. WHO. WHO Director-General declares mpox outbreak a public health emergency of international concern. https://www.who.int/news/item/14-08-2024-who-director-general-declares-mpox-outbreak-a-public-health-emergency-of-international-concern (2024).

18. Kinganda-Lusamaki, E. et al. Clade I mpox virus genomic diversity in the Democratic Republic of the Congo, 2018-2024: Predominance of zoonotic transmission. Cell 188, 4–14.e6 (2025).

19. Khodakevich, L., Jezek, Z. & Kinzanzka, K. Isolation of monkeypox virus from wild squirrel infected in nature. Lancet 1, 98–99 (1986).

20. Leendertz, F., et al. Fire-footed rope squirrels (Funisciurus pyrropus) are a reservoir host of monkeypox virus (Orthopoxvirus monkeypox). Preprint at 10.21203/rs.3.rs-6322223/v1 (2025).

21. Curaudeau, M. et al. Identifying the Most Probable Mammal Reservoir Hosts for Monkeypox Virus Based on Ecological Niche Comparisons. Viruses 15, 727 (2023).

22. Khodakevich, L. et al. Monkeypox virus in relation to the ecological features surrounding human settlements in Bumba zone, Zaire. Trop Geogr Med 39, 56–63 (1987).

23. Hutin, Y. J. et al. Outbreak of human monkeypox, Democratic Republic of Congo, 1996 to 1997. Emerg Infect Dis 7, 434–438 (2001).

24. Reynolds, M. G. et al. A Silent Enzootic of an Orthopoxvirus in Ghana, West Africa: Evidence for Multi-Species Involvement in the Absence of Widespread Human Disease. Am J Trop Med Hyg 82, 746–754 (2010).

25. Doty, J. B. et al. Assessing Monkeypox Virus Prevalence in Small Mammals at the Human-Animal Interface in the Democratic Republic of the Congo. Viruses 9, (2017).

26. Doshi, R. H. et al. Epidemiologic and Ecologic Investigations of Monkeypox, Likouala Department, Republic of the Congo, 2017. Emerg Infect Dis 25, 273–281 (2019).

27. Radonić, A. et al. Fatal monkeypox in wild-living sooty mangabey, Côte d’Ivoire, 2012. Emerg Infect Dis 20, 1009–1011 (2014).

28. Patrono, L. V. et al. Monkeypox virus emergence in wild chimpanzees reveals distinct clinical outcomes and viral diversity. Nat Microbiol 5, 955–965 (2020).

29. Brien, S. C. et al. Clinical Manifestations of an Outbreak of Monkeypox Virus in Captive Chimpanzees in Cameroon, 2016. The Journal of Infectious Diseases 229, S275–S284 (2024).

30. Parker, E. et al. Genomics reveals zoonotic and sustained human mpox spread in West Africa. Nature 643, 1343–1351 (2025).

31. Tiee, M. S., Harrigan, R. J., Thomassen, H. A. & Smith, T. B. Ghosts of infections past: using archival samples to understand a century of monkeypox virus prevalence among host communities across space and time. R Soc Open Sci 5, 171089 (2018).

32. Falendysz, E. A. et al. Characterization of Monkeypox virus infection in African rope squirrels (Funisciurus sp.). PLoS Negl Trop Dis 11, e0005809 (2017).

33. Amman, B. R. et al. Seasonal pulses of Marburg virus circulation in juvenile Rousettus aegyptiacus bats coincide with periods of increased risk of human infection. PLoS Pathogens 8, (2012).

34. Meta Djomsi, D., et al. Coronaviruses Are Abundant and Genetically Diverse in West and Central African Bats, including Viruses Closely Related to Human Coronaviruses. Viruses 15, 337 (2023).

35. Lourie, B., Nakano, J. H., Kemp, G. E. & Setzer, H. W. Isolation of poxvirus from an African Rodent. J Infect Dis 132, 677–681 (1975).

36. Kingdon, J. The Kingdon Field Guide to African Mammals: Second *Edition*. (Bloomsbury Wildlife (Bloomsbury Naturalist series), London, 2018).

37. Irwin, D. M., Kocher, T. D. & Wilson, A. C. Evolution of the cytochromeb gene of mammals. J Mol Evol 32, 128–144 (1991).

38. Montgelard, C., Bentz, S., Tirard, C., Verneau, O. & Catzeflis, F. M. Molecular systematics of sciurognathi (rodentia): the mitochondrial cytochrome b and 12S rRNA genes support the Anomaluroidea (Pedetidae and Anomaluridae). Mol Phylogenet Evol 22, 220–233 (2002).

39. Lacroix, A. et al. Investigating the Circulation of Ebola Viruses in Bats during the Ebola Virus Disease Outbreaks in the Equateur and North Kivu Provinces of the Democratic Republic of Congo from 2018. Pathogens 10, 557 (2021).

40. Kocher, T. D. et al. Dynamics of mitochondrial DNA evolution in animals: amplification and sequencing with conserved primers. Proc. Natl. Acad. Sci. U.S.A. 86, 6196– 6200 (1989).

41. Gouy, M., Guindon, S. & Gascuel, O. SeaView Version 4: A Multiplatform Graphical User Interface for Sequence Alignment and Phylogenetic Tree Building. Mol Biol Evol 27, 221– 224 (2010).

42. Schroeder, K. & Nitsche, A. Multicolour, multiplex real-time PCR assay for the detection of human-pathogenic poxviruses. Molecular and Cellular Probes 24, 110–113 (2010).

43. Katoh, K. & Standley, D. M. MAFFT Multiple Sequence Alignment Software Version 7: Improvements in Performance and Usability. Molecular Biology and Evolution 30, 772–780 (2013).

44. Kalyaanamoorthy, S., Minh, B. Q., Wong, T. K. F., von Haeseler, A. & Jermiin, L. S. ModelFinder: fast model selection for accurate phylogenetic estimates. Nat Methods 14, 587– 589 (2017).

45. Minh, B. Q. et al. IQ-TREE 2: New Models and Efficient Methods for Phylogenetic Inference in the Genomic Era. Mol Biol Evol 37, 1530–1534 (2020).

46. Hoang, D. T., Chernomor, O., von Haeseler, A., Minh, B. Q. & Vinh, L. S. UFBoot2: Improving the Ultrafast Bootstrap Approximation. Mol Biol Evol 35, 518–522 (2018).

47. Hadfield, J. et al. Nextstrain: real-time tracking of pathogen evolution. Bioinformatics 34, 4121–4123 (2018).

48. Kinganda-Lusamaki, E. et al. Use of Mpox Multiplex Serology in the Identification of Cases and Outbreak Investigations in the Democratic Republic of the Congo (DRC). Pathogens 12, 916 (2023).

49. Dubois, M. E., Hammarlund, E. & Slifka, M. K. Optimization of Peptide-Based ELISA for Serological Diagnostics: A Retrospective Study of Human Monkeypox Infection. Vector-Borne and Zoonotic Diseases 12, 400–409 (2012).

50. Ayouba, A. et al. Development of a Sensitive and Specific Serological Assay Based on Luminex Technology for Detection of Antibodies to Zaire Ebola Virus. Journal of Clinical Microbiology 55, 165–176 (2017).

51. Peeters, M. et al. Extensive Survey and Analysis of Factors Associated with Presence of Antibodies to Orthoebolaviruses in Bats from West and Central Africa. Viruses 15, 1927 (2023).

