## Supplementary Table S1 for "High genetic diversity of mpox virus (MPXV) in three different rodent species in the Democratic Republic of the Congo (DRC)"

**Supplementary table 1.** Animals sampled and tested by PCR for the presence of Orthopoxviruses per province and collection site, by animal type, genus and species. Provinces correspond to those shown in Fig. 1.

| Animal type/species | Bas Uele | | | | | Total Bas Uele | Equateur | | | | Total Equateur | Ituri | Kinshasa | Kwilu | Sud-Kivu | | Total Sud-Kivu | Sud-Ubangi | Tshopo | | | | | | | | | | | | | | | Total Tshopo | Tshuapa | TOTAL |
|---|---|---|---|---|---|---|---|---|---|---|---|---|---|---|---|---|---|---|---|---|---|---|---|---|---|---|---|---|---|---|---|---|---|---|---|---|
| | BBO | BOG | KPN | LIK | WEL | | BIK | BLB | ING | MBK | | EPL | KZA | KIK | KMB | LWK | | GEM | BOM | BWI | HMB | KIS | KON | LEK | MBG | MLK | MSK | OBE | UMA | WKO | YAI | YKO | YLG | | INK | |
| **O/ Chiroptera** | | | **103** | | | **103** | **79** | | | **20** | **99** | | | | **50** | **49** | **99** | **101** | | **71** | | **35** | | | | **17** | **25** | | | **17** | | | **11** | **176** | **136** | **714** |
| **SO/Yinpterochiroptera** | | | **44** | | | **44** | **77** | | | **18** | **95** | | | | **50** | **48** | **98** | **101** | | **71** | | **29** | | | | **16** | **25** | | | **15** | | | **10** | **166** | **132** | **636** |
| *Casinycteris argynnis* | | | 4 | | | 4 | | | | | | | | | | | | | | 2 | | | | | | 2 | | | | | | | | 4 | 7 | 15 |
| *Doryrhina camerunensis* | | | | | | | | | | | | | | | | | | | | | | | | | | | | | | | | | | | 3 | 3 |
| *Doryrhina cyclops* | | | 3 | | | 3 | | | | | | | | | | | | | | | | | | | | | | | | | | | | | | 3 |
| *Eidolon helvum* | | | | | | | 1 | | | | 1 | | | | | | | 52 | | | | | | | | | | | | | | | | | 27 | 80 |
| *Epomophorus* sp. | | | | | | | 69 | | | 18 | 87 | | | | 43 | 43 | 86 | | | | | | | | | | | | | | | | | | | 173 |
| *Epomops franqueti* | | | 1 | | | 1 | 6 | | | | 6 | | | | | | | 26 | | 43 | | 1 | | | | 12 | 17 | | | 10 | | | 1 | 84 | 20 | 137 |
| *Hipposideros beatus* | | | 2 | | | 2 | 1 | | | | 1 | | | | | | | | | | | | | | | | | | | | | | 1 | 1 | 5 | 9 |
| *Hipposideros fuliginosus* | | | 4 | | | 4 | | | | | | | | | | | | | | | | | | | | | | | | | | | | | | 4 |
| *Hipposideros gigas* | | | 1 | | | 1 | | | | | | | | | | | | | | | | 1 | | | | | | | | | | | | 1 | | 2 |
| *Hipposideros ruber* | | | 1 | | | 1 | | | | | | | | | | | | | | | | | | | | | | | | | | | | | | 1 |
| *Hipposideros* sp. | | | 1 | | | 1 | | | | | | | | | | | | | | | | | | | | | | | | | | | | | | 1 |
| *Hypsignathus monstrosus* | | | 3 | | | 3 | | | | | | | | | | | | 15 | | 2 | | | | | | | | | | | | | | 2 | 11 | 31 |
| *Lavia frons* | | | | | | | | | | | | | | | | | | | | | | | | | | 1 | | | | | | | 1 | 2 | | 2 |
| *Lissonycteris angolensis* | | | | | | | | | | | | | | | | 3 | 3 | | | | | | | | | | | | | | | | | | | 3 |
| *Megaloglossus woermanni* | | | 3 | | | 3 | | | | | | | | | 1 | | 1 | 3 | | 14 | | 21 | | | | 1 | 5 | | | | | | | 41 | 21 | 69 |
| *Myonycteris* sp. | | | | | | | | | | | | | | | | | | | | | | | | | | | | | | | | | | | 16 | 16 |
| *Myonycteris torquata* | | | 5 | | | 5 | | | | | | | | | | | | 5 | | 10 | | 6 | | | | | 3 | | | 5 | | | 7 | 31 | 18 | 59 |
| *Rhinolophus alcyone* | | | 3 | | | 3 | | | | | | | | | | | | | | | | | | | | | | | | | | | | | | 3 |
| *Rousettus aegyptiacus* | | | | | | | | | | | | | | | 2 | 2 | 4 | | | | | | | | | | | | | | | | | | | 4 |
| *Scotonycteris bergmansi* | | | 2 | | | 2 | | | | | | | | | | | | | | | | | | | | | | | | | | | | | 1 | 3 |
| *Stenonycteris lanosa* | | | | | | | | | | | | | | | 4 | | 4 | | | | | | | | | | | | | | | | | | | 4 |
| *Pteropodidae NA* | | | 11 | | | 11 | | | | | | | | | | | | | | | | | | | | | | | | | | | | | 3 | 14 |
| **SO/Yangochiroptera** | | | **59** | | | **59** | **2** | | | **2** | **4** | | | | | **1** | **1** | | | | | **6** | | | | **1** | | | | **2** | | | **1** | **10** | **4** | **78** |
| *Afronycteris nanus* | | | 2 | | | 2 | | | | | | | | | | | | | | | | 3 | | | | | | | | | | | 1 | 4 | 1 | 7 |
| *Glauconycteris alboguttata* | | | 1 | | | 1 | | | | | | | | | | | | | | | | | | | | | | | | | | | | | | 1 |
| *Glauconycteris beatrix* | | | 1 | | | 1 | | | | | | | | | | | | | | | | | | | | | | | | | | | | | 1 | 2 |
| *Glauconycteris cf. humeralis* | | | 4 | | | 4 | | | | | | | | | | | | | | | | | | | | | | | | | | | | | | 4 |
| *Glauconycteris curryae* | | | 4 | | | 4 | | | | | | | | | | | | | | | | | | | | | | | | | | | | | | 4 |
| *Glauconycteris* sp. | | | | | | | | | | | | | | | | | | | | | | | | | | | | | | 1 | | | | 1 | 1 | 2 |
| *Glauconycteris variegata* | | | 1 | | | 1 | | | | | | | | | | | | | | | | | | | | | | | | | | | | | | 1 |
| *Kerivoula cuprosa* | | | 1 | | | 1 | | | | | | | | | | | | | | | | | | | | | | | | | | | | | | 1 |
| *Kerivoula lanosa* | | | | | | | | | | | | | | | | | | | | | | | | | | | | | | | | | | | 1 | 1 |
| *Kerivoula* sp. | | | 1 | | | 1 | | | | | | | | | | | | | | | | | | | | | | | | | | | | | | 1 |
| *Mimetillus moloneyi* | | | 2 | | | 2 | | | | | | | | | | | | | | | | | | | | 1 | | | | | | | | 1 | | 3 |
| *Mops* sp. | | | 28 | | | 28 | 2 | | | 1 | 3 | | | | | 1 | 1 | | | | | | | | | | | | | | | | | | | 32 |
| *Neoromicia rendalli* | | | 1 | | | 1 | | | | | | | | | | | | | | | | | | | | | | | | | | | | | | 1 |
| *Nycticeinops happoldorum* | | | 3 | | | 3 | | | | | | | | | | | | | | | | | | | | | | | | | | | | | | 3 |
| *Nycticeinops* sp. | | | | | | | | | | | | | | | | | | | | | | | | | | | | | | 1 | | | | 1 | | 1 |
| *Pipistrellus nanulus* | | | 5 | | | 5 | | | | | | | | | | | | | | | | | | | | | | | | | | | | | | 5 |
| *Pseudoromicia mbamminkom* | | | 2 | | | 2 | | | | | | | | | | | | | | | | | | | | | | | | | | | | | | 2 |
| *Scotophilus nux* | | | 2 | | | 2 | | | | | | | | | | | | | | | | | | | | | | | | | | | | | | 2 |
| *Scotophilus* sp. | | | | | | | | | | 1 | 1 | | | | | | | | | | | | | | | | | | | | | | | | | 1 |
| *Vansonia/Pipistrellus rueppellii* | | | | | | | | | | | | | | | | | | | | | | 3 | | | | | | | | | | | | 3 | | 3 |
| *Vespertilionidae NA* | | | 1 | | | 1 | | | | | | | | | | | | | | | | | | | | | | | | | | | | | | 1 |
| **O/Rodentia** | | | | | | | | | | | | | | | | | | | | | | | | | | | | | | | | | | | | |
| **SO/ Myomoprha** | | | **273** | | | **273** | **41** | **8** | **46** | | **95** | | **2** | | **46** | **33** | **79** | **19** | | **41** | | **52** | | | | **20** | **97** | | | **54** | | | **32** | **296** | **262** | **1026** |
| *Aethomys* sp. | | | | | | | | | | | | | 1 | | | | | | | | | | | | | | | | | | | | | | | 1 |
| *Congomys lukolelae* | | | | | | | | | | | | | | | | | | | | | | | | | | | | | | | | | | | 3 | 3 |
| *Cricetomys gambianus* | | | | | | | 5 | | | | 5 | | | | | | | | | | | | | | | | | | | | | | | | | 5 |
| *Cricetomys* sp. | | | | | | | | | | | | | | | | | | | | | | | | | | | | | | | | | | | 1 | 1 |
| *Dasymys* sp. | | | | | | | | | | | | | | | 1 | | 1 | | | | | | | | | | | | | | | | | | | 1 |
| *Dendromus* sp. | | | | | | | 1 | | | | 1 | | 1 | | | | | | | 2 | | | | | | | | | | | | | 1 | 3 | 5 | 10 |
| *Deomys ferrugineus* | | | | | | | | | | | | | | | | | | | | | | | | | | 1 | 4 | | | | | | 4 | 9 | | 9 |
| *Grammomys poensis* | | | 1 | | | 1 | | | | | | | | | | | | | | | | | | | | | | | | | | | | | | 1 |
| *Grammomys* sp. | | | 2 | | | 2 | | | | | | | | | | | | | | | | | | | | | | | | 3 | | | | 3 | | 5 |
| *Hybomys lunaris* | | | 9 | | | 9 | | | | | | | | | | | | | | | | | | | | | | | | | | | 2 | 2 | 5 | 16 |
| *Hybomys* sp. | | | 14 | | | 14 | 1 | 7 | 13 | | 21 | | | | | | | | | | | | | | | | 11 | | | 1 | | | | 12 | | 47 |
| *Hylomyscus aeta* | | | 2 | | | 2 | | | | | | | | | | | | | | | | | | | | | | | | | | | | | | 2 |
| *Hylomyscus* sp. | | | | | | | | 1 | 5 | | 6 | | | | | | | | | | | | | | | | | | | | | | | | 30 | 36 |
| *Hylomyscus stella* | | | 14 | | | 14 | | | | | | | | | | | | | | | | | | | | | 2 | | | 2 | | | | 4 | | 18 |
| *Hylomyscus thornesmithae* | | | | | | | | | | | | | | | | | | | | | | | | | | | | | | | | | | | 9 | 9 |
| *Hylomyscus alleni* | | | 13 | | | 13 | | | | | | | | | | | | 2 | | | | | | | | | | | | | | | | | | 15 |
| *Lemniscomys* sp. | | | 1 | | | 1 | | | | | | | | | | | | | | | | | | | | | | | | | | | | | | 1 |

| Animal type/species | Bas Uele | | | | | Total Bas Uele | Equateur | | | | Total Equateur | Ituri | Kinshasa | Kwilu | Sud-Kivu | | Total Sud-Kivu | Sud-Ubangi | Tshopo | | | | | | | | | | | | | | | Total Tshopo | Tshuapa | TOTAL |
|---|---|---|---|---|---|---|---|---|---|---|---|---|---|---|---|---|---|---|---|---|---|---|---|---|---|---|---|---|---|---|---|---|---|---|---|---|
| | BBO | BOG | KPN | LIK | WEL | | BIK | BLB | ING | MBK | | EPL | KZA | KIK | KMB | LWK | | GEM | BOM | BWI | HMB | KIS | KON | LEK | MBG | MLK | MSK | OBE | UMA | WKO | YAI | YKO | YLG | | INK | |
| *Lemniscomys striatus* | | | 2 | | | 2 | 5 | | | | 5 | | | | 2 | | 2 | | | | | 1 | | | | 5 | 3 | | | 1 | | | 7 | 17 | 5 | 31 |
| *Lophuromys cf. cinereus* | | | | | | | | | | | | | | | 3 | 3 | 6 | | | | | | | | | | | | | | | | | | | 6 |
| *Lophuromys dudui* | | | 20 | | | 20 | | | | | | | | | | | | | | 12 | | 3 | | | | 6 | 36 | | | 8 | | | 9 | 74 | | 94 |
| *Lophuromys huttereri* | | | | | | | | | | | | | | | | | | | | | | | | | | | | | | | | | | | 6 | 6 |
| *Lophuromys rita* | | | | | | | | | | | | | | | | | | | | | | | | | | | | | | | | | | | 38 | 38 |
| *Lophuromys* sp. | | | 2 | | | 2 | 1 | | | | 1 | | | | | | | | | | | 2 | | | | | 1 | | | | | | | 3 | | 6 |
| *Malacomys longipes* | | | 5 | | | 5 | | | | | | | | | | | | | | 1 | | | | | | | | | | 1 | | | | 2 | 3 | 10 |
| *Mastomys natalensis* | | | 9 | | | 9 | | | | | | | | | | | | 15 | | 10 | | 1 | | | | | 7 | | | | | | 2 | 20 | | 44 |
| *Mastomys* sp. | | | | | | | | | | | | | | | 2 | 1 | 3 | | | | | | | | | | | | | | | | | | | 3 |
| *Mus cf. gratus* | | | 16 | | | 16 | | | | | | | | | | | | | | | | | | | | | | | | | | | | | 31 | 47 |
| *Mus setulosus* | | | 1 | | | 1 | | | | | | | | | | | | | | | | | | | | | | | | | | | | | | 1 |
| *Mus* sp. | | | 5 | | | 5 | 8 | | | | 8 | | | | 6 | 2 | 8 | 1 | | 12 | | | | | | 2 | 3 | | | 3 | | | 1 | 21 | | 43 |
| *Oenomys hypoxanthus* | | | 3 | | | 3 | | | | | | | | | | | | | | | | | | | | | 2 | | | | | | 2 | 4 | 1 | 8 |
| *Oenomys* sp. | | | | | | | | | 11 | | 11 | | | | | | | | | | | | | | | | | | | | | | | | | 11 |
| *Pelomys* sp. | | | | | | | | | | | | | | | 1 | | 1 | | | | | | | | | | | | | | | | | | | 1 |
| *Praomys jacksoni* | | | 128 | | | 128 | 10 | | | | 10 | | | | | | | 1 | | 1 | | 10 | | | | 5 | 21 | | | 28 | | | 1 | 66 | | 205 |
| *Praomys minor* | | | | | | | | | | | | | | | | | | | | | | 13 | | | | | | | | | | | | 13 | 31 | 44 |
| *Praomys misonnei* | | | 19 | | | 19 | | | | | | | | | | | | | | | | | | | | | | | | | | | | | | 19 |
| *Praomys mutoni* | | | | | | | | | | | | | | | | | | | | | | | | | | | | | | | | | | | 5 | 5 |
| *Praomys* sp. | | | 3 | | | 3 | | | 2 | | 2 | | | | | | | | | | | | | | | | | | | | | | | | 4 | 9 |
| *Praomys verschureni* | | | 1 | | | 1 | | | | | | | | | | | | | | | | | | | | | | | | | | | | | | 1 |
| *Rattus norvegicus* | | | | | | | | | | | | | | | | | | | | | | 12 | | | | | | | | | | | | 12 | | 12 |
| *Rattus rattus* | | | 1 | | | 1 | 10 | | 15 | | 25 | | | | 30 | 25 | 55 | | | 3 | | 10 | | | | 1 | 7 | | | | | | 3 | 24 | 72 | 177 |
| *Stochomys longicaudatus* | | | | | | | | | | | | | | | | | | | | | | | | | | | | | | | | | | | 9 | 9 |
| *Tachyoryctes* sp. | | | | | | | | | | | | | | | | 2 | 2 | | | | | | | | | | | | | | | | | | | 2 |
| *Thamnomys poensis* | | | | | | | | | | | | | | | | | | | | | | | | | | | | | | | | | | | 4 | 4 |
| *Thamnomys* sp. | | | 2 | | | 2 | | | | | | | | | | | | | | | | | | | | | | | | 7 | | | | 7 | | 9 |
| *Zelotomys hildegardeae* | | | | | | | | | | | | | | | 1 | | 1 | | | | | | | | | | | | | | | | | | | 1 |
| **SO/ Sciuromorpha** | **19** | **6** | **23** | **10** | **13** | **71** | **21** | | | | **21** | **43** | | **16** | | | | **6** | **3** | | **5** | | **2** | **4** | **4** | **1** | | **5** | **18** | **75** | **3** | **289** | **5** | **414** | **16** | **587** |
| *Funisciurus anerythrus* | 12 | 4 | | 7 | 8 | 31 | | | | | | 33 | | | | | | | | | 1 | | | | 2 | | | 4 | 13 | | 1 | 193 | | 214 | | 278 |
| *Funisciurus cfr bayonii* | | | | | | | | | | | | | | | | | | | | | | | | | | | | | | | | 72 | | 72 | | 72 |
| *Funisciurus congolensis* | | | | | | | | | | | | | | 15 | | | | | | | | | | | | | | | | | | | | | | 15 |
| *Funisciurus* sp. | 1 | 2 | 20 | 1 | 2 | 26 | 16 | | | | 16 | 2 | | | | | | 6 | 3 | | 1 | | 1 | 4 | 2 | 1 | | | 3 | 39 | 1 | 1 | 1 | 57 | 9 | 116 |
| *Heliosciurus rufobrachium* | 2 | | | | | 2 | | | | | | | | | | | | | | | | | | | | | | | | | | | | | 1 | 3 |
| *Heliosciurus* sp. | | | 2 | | | 2 | 5 | | | | 5 | | | | | | | | | | 3 | | | | | | | | | 1 | | 1 | | 5 | | 12 |
| *Paraxerus boehmi* | 1 | | | 2 | | 3 | | | | | | 1 | | | | | | | | | | | | | | | | | | | | 1 | | 1 | | 5 |
| *Paraxerus cfr alexandri* | | | | | | | | | | | | 1 | | | | | | | | | | | | | | | | | | | | | | | | 1 |
| *Paraxerus* sp. | 1 | | 1 | | 2 | 4 | | | | | | 6 | | | | | | | | | | | | | | | | 1 | 1 | 33 | 1 | 3 | 4 | 43 | | 53 |
| *Protoxerus* sp. | 1 | | | | | 1 | | | | | | | | 1 | | | | | | | | | | | | | | | | 2 | | 9 | | 11 | | 13 |
| *Protoxerus stangeri* | 1 | | | | | 1 | | | | | | | | | | | | | | | | | 1 | | | | | | | | | 2 | | 3 | | 4 |
| *NA* | | | | | 1 | 1 | | | | | | | | | | | | | | | | | | | | | | | 1 | | | 7 | | 8 | 6 | 15 |
| **O/ Soricomorpha** | | | **46** | | | **46** | **9** | | | **3** | **12** | | | | **4** | **17** | **21** | **1** | | **10** | | **21** | | | | **11** | **25** | | | **36** | | | **14** | **117** | **130** | **327** |
| *Crocidura caliginea* | | | 3 | | | 3 | | | | | | | | | | | | | | | | | | | | | 1 | | | | | | | 1 | | 4 |
| *Crocidura cf. denti* | | | 8 | | | 8 | | | | | | | | | | | | | | | | 8 | | | | 7 | 8 | | | 11 | | | 5 | 39 | | 47 |
| *Crocidura crenata* | | | 1 | | | 1 | | | | | | | | | | | | | | | | | | | | | | | | | | | | | | 1 |
| *Crocidura dolichura* | | | 1 | | | 1 | | | | | | | | | | | | | | | | | | | | | 1 | | | | | | | 1 | | 2 |
| *Crocidura grassei* | | | | | | | | | | | | | | | | | | | | | | | | | | | | | | | | | | | 1 | 1 |
| *Crocidura littoralis* | | | 1 | | | 1 | | | | | | | | | | | | | | 5 | | | | | | 1 | 7 | | | 17 | | | 6 | 36 | | 37 |
| *Crocidura ludia* | | | 1 | | | 1 | | | | | | | | | | | | | | 2 | | | | | | 2 | | | | 4 | | | 1 | 9 | | 10 |
| *Crocidura olivieri* | | | 17 | | | 17 | 9 | | | 3 | 12 | | | | 3 | 15 | 18 | 1 | | 2 | | 8 | | | | 1 | 6 | | | 2 | | | 1 | 20 | 16 | 84 |
| *Crocidura* sp. | | | 1 | | | 1 | | | | | | | | | 1 | 2 | 3 | | | 1 | | 5 | | | | | | | | | | | | 6 | 91 | 101 |
| *Paracrocidura cf. maxima* | | | | | | | | | | | | | | | | | | | | | | | | | | | | | | | | | | | 4 | 4 |
| *Paracrocidura schoutedeni* | | | 3 | | | 3 | | | | | | | | | | | | | | | | | | | | | | | | | | | | | | 3 |
| *Paracrocidura* sp. | | | | | | | | | | | | | | | | | | | | | | | | | | | | | | 1 | | | | 1 | | 1 |
| *Scutisorex congicus* | | | 6 | | | 6 | | | | | | | | | | | | | | | | | | | | | | | | | | | 1 | 1 | | 7 |
| *Scutisorex* sp. | | | | | | | | | | | | | | | | | | | | | | | | | | | 2 | | | | | | | 2 | 6 | 8 |
| *Sylvisorex johnstoni* | | | 4 | | | 4 | | | | | | | | | | | | | | | | | | | | | | | | | | | | | | 4 |
| *Sylvisorex* sp. | | | | | | | | | | | | | | | | | | | | | | | | | | | | | | 1 | | | | 1 | 12 | 13 |
| **Other mammals** | | | | | | | **4** | | | | **4** | | | | | | | **1** | | | | | | | | **1** | | | | | | | **1** | **2** | **40** | **47** |
| *Cercopithecus ascanius* | | | | | | | | | | | | | | | | | | 1 | | | | | | | | | | | | | | | | | | 1 |
| *Chrysochloris stuhlmanni* | | | | | | | | | | | | | | | | | | | | | | | | | | 1 | | | | | | | | 1 | | 1 |
| *Galagoides demidoff* | | | | | | | 2 | | | | 2 | | | | | | | | | | | | | | | | | | | | | | | | 1 | 3 |
| *Galagoides thomasi* | | | | | | | | | | | | | | | | | | | | | | | | | | | | | | | | | 1 | 1 | | 1 |
| *Petrodromus tetradactylus* | | | | | | | 2 | | | | 2 | | | | | | | | | | | | | | | | | | | | | | | | 39 | 41 |
| **TOTAL** | **19** | **6** | **445** | **10** | **13** | **493** | **154** | **8** | **46** | **23** | **231** | **43** | **2** | **16** | **100** | **99** | **199** | **128** | **3** | **122** | **5** | **108** | **2** | **4** | **4** | **50** | **147** | **5** | **18** | **182** | **3** | **289** | **63** | **1005** | **584** | **2701** |
