## Supplementary Table S2 for "High genetic diversity of mpox virus (MPXV) in three different rodent species in the Democratic Republic of the Congo (DRC)"

**Supplementary Table 2.** Number of samples (N) analyzed by PCR for the presence of Orthopoxvirus by sample type. Sample types are listed using two-letter abbreviation, and organ pools are designated from pool 1 to 4. Sample number testing positive for OPV PCR are into brackets. Tested organs tested are identified as follow: SP: Spleen, LI: Liver, LU: Lung, KI: Kidney, FE: Faeces, IN: Intestine, SA: Oral swab, RE: Rectal swab, TE: Testis, TO: Tongue, EY: Eye, BR: Brain. Pools contain the following organs: Pool 1: SP+LU; Pool 2: LI+KI; Pool 3: SP+LI+LU+KI+TE+TO and Pool 4: TE+BR.

| Animal Order /SubOrder | No. sampled animals * | No. SP | No. LI | No. LU | No. KI | No. FE | No. IN | No. SA | No. RE | No. TE | No. TO | No. EY | No. BR | No. Pool 1 | No. Pool 2 | No. Pool 3 | No. Pool 4 | Total No. of samples tested |
|---|---|---|---|---|---|---|---|---|---|---|---|---|---|---|---|---|---|---|
| **O/ Chiroptera** | | | | | | | | | | | | | | | | | | |
| SO/Yinpterochiroptera | 636 | 312 | 3 | 16 | - | - | 1 | - | 98 | - | - | - | - | 35 | 182 | - | - | 647 |
| SO/Yangochiroptera | 78 | 9 | 2 | 2 | 1 | - | - | - | 1 | - | - | - | - | 1 | 63 | - | - | 79 |
| **O/ Rodentia** | | | | | | | | | | | | | | | | | | |
| SO/Myomorpha | 1027 | 371 | 1 | 71 | 15 | 13 | - | 54 | 54 | 13 | 12 | 12 | - | 101 (1) | 550 | - | - | 1267 (1) |
| SO/ Sciuromorpha | 586 | 32 (1) | 440 (2) | 4 | 2 | 2 | - | - | - | 1 | 2 | | - | 62 | 39 | 12 | - | 596 (3) |
| **O/ Soricomorpha** | 327 | 109 (1) | 2 | 51 | 9 | 10 | - | - | - | 9 | 10 | | 1 | 31 | 182 | 1(1) | 1 | 416 (2) |
| **Other mammals[£]** | 47 | 6 | - | - | - | - | - | - | - | - | - | - | - | 1 | 40 | - | - | 47 |
| **TOTAL** | **2701** | **839 (2)** | **448 (2)** | **144** | **27** | **25** | **1** | **54** | **153** | **23** | **24** | **12** | **1** | **231 (1)** | **1056** | **13 (1)** | **1** | **3052 (6)** |
