## Supplementary Table S3 for "High genetic diversity of mpox virus (MPXV) in three different rodent species in the Democratic Republic of the Congo (DRC)"

**Supplementary table 3.** Number of animals tested for presence of antibodies against MPXV antigen by Luminex serology, by animal species and order.

| | Total tested | animal positive to at least 1 Ag | animal positive to 1 Ag | animal positive to 2 Ag | animal negative | Positive Ag B21R 179/180 | Positive Ag B21R 180/186 | Positive Ag B22R 64/65 | Overall total |
|---|---|---|---|---|---|---|---|---|---|
| **O/Chiroptera** | **633** | **14** | **12** | **2** | **619** | **6** | **4** | **6** | **633** |
| **SO/Yinpterochiroptera** | **563** | **13** | **11** | **2** | **550** | **5** | **4** | **6** | **563** |
| *Casinycteris argynnis* | 13 | | | | 13 | | | | 13 |
| *Doryrhina camerunensis* | 3 | | | | 3 | | | | 3 |
| *Doryrhina cyclops* | 3 | | | | 3 | | | | 3 |
| *Eidolon helvum* | 80 | 6 | 5 | 1 | 74 | 1 | 1 | 5 | 80 |
| *Epomophorus* sp. | 161 | 6 | 5 | 1 | 155 | 4 | 2 | 1 | 161 |
| *Epomops franqueti* | 98 | 1 | 1 | | 97 | | 1 | | 98 |
| *Hipposideros beatus* | 8 | | | | 8 | | | | 8 |
| *Hipposideros fuliginosus* | 4 | | | | 4 | | | | 4 |
| *Hipposideros gigas* | 2 | | | | 2 | | | | 2 |
| *Hipposideros ruber* | 1 | | | | 1 | | | | 1 |
| *Hipposideros* sp. | 1 | | | | 1 | | | | 1 |
| *Hypsignathus monstrosus* | 31 | | | | 31 | | | | 31 |
| *Lavia frons* | 1 | | | | 1 | | | | 1 |
| *Lissonycteris angolensis* | 3 | | | | 3 | | | | 3 |
| *Megaloglossus woermanni* | 60 | | | | 60 | | | | 60 |
| *Myonycteris torquata* | 50 | | | | 16 | | | | 50 |
| *Myonycteris* sp. | 16 | | | | 50 | | | | 16 |
| *Rhinolophus alcyone* | 3 | | | | 3 | | | | 3 |
| *Rousettus aegyptiacus* | 4 | | | | 14 | | | | 4 |
| *Scotonycteris bergmansi* | 3 | | | | 4 | | | | 3 |
| *Stenonycteris lanosa* | 4 | | | | 3 | | | | 4 |
| *Pteropodidae NA* | 14 | | | | 4 | | | | 14 |
| **SO/Yangochiroptera** | **70** | **1** | **1** | | **69** | **1** | | | **70** |
| *Afronycteris nanus* | 5 | | | | 5 | | | | 5 |
| *Glauconycteris alboguttata* | 1 | | | | 1 | | | | 1 |
| *Glauconycteris beatrix* | 2 | | | | 2 | | | | 2 |
| *Glauconycteris cf. humeralis* | 4 | | | | 4 | | | | 4 |
| *Glauconycteris curryae* | 4 | | | | 4 | | | | 4 |
| *Glauconycteris* sp. | 1 | | | | 1 | | | | 1 |
| *Glauconycteris variegata* | 1 | | | | 1 | | | | 1 |
| *Kerivoula cuprosa* | 1 | | | | 1 | | | | 1 |
| *Kerivoula lanosa* | 1 | | | | 1 | | | | 1 |
| *Kerivoula* sp. | 1 | | | | 1 | | | | 1 |
| *Mimetillus moloneyi* | 2 | | | | 2 | | | | 2 |
| *Mops* sp. | 32 | 1 | 1 | | 31 | 1 | | | 32 |
| *Neoromicia rendalli* | 1 | | | | 1 | | | | 1 |
| *Nycticeinops happoldorum* | 3 | | | | 3 | | | | 3 |
| *Pipistrellus nanulus* | 5 | | | | 5 | | | | 5 |
| *Pseudoromicia mbamminkom* | 2 | | | | 2 | | | | 2 |
| *Scotophilus nux* | 2 | | | | 2 | | | | 2 |
| *Vansonia/Pipistrellus rueppellii* | 1 | | | | 1 | | | | 1 |
| *Vespertilionidae NA* | 1 | | | | 1 | | | | 1 |
| **O/Rodentia** | **723** | **14** | **14** | | **709** | **4** | **7** | **3** | **723** |
| **SO/Myomoprha** | **723** | **14** | **14** | | **709** | **4** | **7** | **3** | **723** |
| *Congomys lukolelae* | 3 | | | | 3 | | | | 3 |
| *Cricetomys* sp. | 1 | | | | 1 | | | | 1 |
| *Dasymys* sp. | 1 | | | | 1 | | | | 1 |
| *Dendromus* sp. | 5 | | | | 5 | | | | 5 |
| *Deomys ferrugineus* | 1 | | | | 1 | | | | 1 |
| *Grammomys poensis* | 1 | | | | 1 | | | | 1 |
| *Grammomys* sp. | 2 | | | | 2 | | | | 2 |
| *Hybomys lunaris* | 13 | | | | 13 | | | | 13 |
| *Hybomys* sp. | 17 | | | | 17 | | | | 17 |
| *Hylomyscus aeta* | 2 | | | | 2 | | | | 2 |
| *Hylomyscus alleni* | 15 | | | | 15 | | | | 15 |
| *Hylomyscus* sp. | 30 | 1 | 1 | | 29 | | 1 | | 30 |
| *Hylomyscus stella* | 16 | | | | 16 | | | | 16 |
| *Hylomyscus thornesmithae* | 9 | | | | 9 | | | | 9 |
| *Lemniscomys* sp. | 1 | | | | 1 | | | | 1 |
| *Lemniscomys striatus* | 10 | | | | 10 | | | | 10 |
| *Lophuromys cf. cinereus* | 6 | | | | 6 | | | | 6 |
| *Lophuromys dudui* | 40 | | | | 40 | | | | 40 |
| *Lophuromys huttereri* | 6 | | | | 6 | | | | 6 |
| *Lophuromys rita* | 38 | | | | 38 | | | | 38 |
| *Lophuromys* sp. | 3 | | | | 3 | | | | 3 |
| *Malacomys longipes* | 9 | | | | 9 | | | | 9 |
| *Mastomys natalensis* | 31 | | | | 31 | | | | 31 |
| *Mastomys* sp. | 3 | | | | 3 | | | | 3 |
| *Mus cf. gratus* | 47 | | | | 47 | | | | 47 |
| *Mus setulosus* | 1 | | | | 1 | | | | 1 |
| *Mus* sp. | 15 | | | | 15 | | | | 15 |
| *Oenomys hypoxanthus* | 5 | | | | 5 | | | | 5 |
| *Pelomys* sp. | 1 | | | | 1 | | | | 1 |
| *Praomys jacksoni* | 141 | 3 | 3 | | 138 | 1 | 1 | 1 | 141 |
| *Praomys minor* | 42 | 3 | 3 | | 39 | | 2 | 1 | 42 |
| *Praomys misonnei* | 19 | 1 | 1 | | 18 | 1 | | | 19 |
| *Praomys mutoni* | 5 | | | | 5 | | | | 5 |
| *Praomys* sp. | 7 | | | | 7 | | | | 7 |
| *Praomys verschureni* | 1 | | | | 1 | | | | 1 |
| *Rattus norvegicus* | 11 | 1 | 1 | | 10 | | | 1 | 11 |
| *Rattus rattus* | 147 | 5 | 5 | | 142 | 2 | 3 | | 147 |
| *Stochomys longicaudatus* | 9 | | | | 9 | | | | 9 |
| *Tachyoryctes* sp. | 2 | | | | 2 | | | | 2 |
| *Thamnomys poensis* | 4 | | | | 4 | | | | 4 |
| *Thamnomys* sp. | 2 | | | | 2 | | | | 2 |
| *Zelotomys hildegardeae* | 1 | | | | 1 | | | | 1 |
| **Overall total** | **1356** | **28** | **26** | **2** | **1328** | **10** | **11** | **9** | **1356** |
